# Causes and consequences of sex-chromosome turnovers in Diptera

**DOI:** 10.1101/2025.10.24.684443

**Authors:** Lorena Layana, Melissa A. Toups, Beatriz Vicoso

## Abstract

Sex-chromosome systems are highly variable across animals, but how they transition from one to another is not well understood. Diptera have undergone multiple sex-chromosome turnovers and expansions while maintaining their general chromosomal content, which makes them an ideal clade to study such transitions. We analysed more than 100 dipteran whole-genome assemblies and identified 4 new lineages that underwent sex-chromosome turnover (in addition to the 5 previously reported). We find the majority of turnovers happened in the group Schizophora, which tend to have fewer genes on the F element (the chromosome homologous to the ancestral insect X chromosome) than lower dipterans, a factor previously hypothesized to facilitate turnover. Most derived X chromosomes have higher GC content than autosomes, consistent with a high prevalence of male-achiasmy in Diptera. In addition, an excess of gene movement out of the X is detected for most of these new X chromosomes, and many of these moved genes have high testis expression in Drosophila, suggesting that out-of-X gene movement contributes to the long-term demasculinization of X chromosomes.

## Introduction

In many organisms, sex is determined by a pair of sex chromosomes (X and Y in the case of male heterogamety, or Z and W in female heterogamety). As sex and separate sexes are ancestral in animals, but sex chromosomes are not, turnover (i.e. the reversal of sex chromosomes to autosomes, and their replacement by a novel sex-chromosome pair) must have occurred repeatedly. Turnover is expected to be particularly common when sex chromosomes are young and/or undifferentiated, as at that point they still maintain most of the features of an autosomal pair (Vicoso 2019). The accumulation of deleterious mutations on a young but already degenerating Y chromosome should favor turnover, as this would yield a new XY pair free of such mutation load (the so-called “hot potato” model of turnover (Blaser et al. 2014)). Indeed, recurrent turnover has been observed in many clades with undifferentiated sex chromosomes, such as several groups of fishes (Ansai et al. 2022; Behrens et al. 2024; El Taher et al.) and frogs (Jeffries et al. 2018; Ma and Veltsos 2021; Evans et al. 2024). On the other hand, once sex chromosomes are fully differentiated, they are hypothesized to become more stable, as reverting a highly specialized X or Y chromosome to an autosome is likely to incur harmful fitness effects (POKORNÁ and KRATOCHVÍL 2009). Y chromosome gene loss should prevent this chromosome from becoming fixed as an autosome. However, fixing the X as an autosome may cause fitness effects in males as a result of misregulation due to relictual dosage compensatory mechanisms or through missing key male-specific genes. An example of the potential stability of well-differentiated X chromosomes is provided by mammals, where the same XY pair has been maintained over 190 million years (Graves 2016). Recent studies have uncovered even older pairs of sex chromosomes, such as those of liverworts (Iwasaki et al. 2021) and cephalopods (Coffing et al. 2025), again supporting the idea that differentiated sex chromosomes may function as an “evolutionary trap”. Alternatively, the long term conservation of some sex chromosomes (by other, currently unknown, factors) may lead to an apparent association with differentiation.

Diptera (flies and mosquitoes) is an ideal order in which to study chromosomal evolution, as in many species these are formed by different combinations of 6 conserved chromosomal arms. These so-called “Muller elements” A to F were first described in *Drosophila*, where their homology was inferred from similar banding patterns of salivary gland chromosomal spreads and similar segregation patterns in their linkage maps (Schaeffer 2018). In all *Drosophila* species, element A is the X chromosome, although additional elements have become sex-linked in many species through fusions of autosomes to the X and/or to the Y (Ellison and Bachtrog 2019). The sequencing of the genomes of both *Drosophila melanogaster* and *Anopheles gambiae* showed that the conservation of gene content of Muller elements extended beyond *Drosophila*, as homology could be detected between the chromosomal arms of the two species based on their gene content (Zdobnov et al. 2002). Recent analyses of dipteran genomes show that the Muller elements of higher Diptera have undergone several rearrangements compared to the ancestral dipteran karyotype (Gries, Jaron and colleagues, submitted), but are well conserved through much of the Brachycera branch (and in particular in the Schizophora section, i.e. muscoid flies and their relatives). Since most of the sex chromosome changes studied previously and in this work occurred in the Schizophora, we describe them in terms of homology to Muller elements.

A chromosome homologous to Element A was also inferred to be the X in *A. gambiae* (Zdobnov et al. 2002), which initially suggested that the X chromosome of dipteran insects might have been another example of a well conserved X chromosome. However, further sampling of the clade showed that element F was instead the ancestral X of the clade, and had been replaced by element A in both *Drosophila* and Anopheles independently (Vicoso and Bachtrog 2015). In *Drosophila*, Muller element F is a highly unusual autosome, which likely reflects its history as a former X chromosome. It carries fewer than 100 genes and is dot-shaped in chromosomal spreads (in *D. melanogaster* is also known as the “dot chromosome”, or as the “fourth chromosome”) (Riddle and Elgin 2018). Element F is almost fully constitutively heterochromatic, and is globally regulated by the protein Painting-of-Fourth (Riddle and Elgin 2018). Similar turnover events were inferred in 6 other lineages, with another 5 having transitioned to undifferentiated sex chromosomes (Vicoso and Bachtrog 2015). Whether the former X chromosome simply reverted to an autosome, or underwent more complex rearrangements, could not be established in the absence of chromosome-level genome assemblies.

This high frequency of turnover of an old and well-differentiated X chromosome is at odds with the evolutionary trap hypothesis, but also with other insects, which typically share an ancient X chromosome homologous to element F (Meisel et al. 2019; Chauhan et al. 2021; Toups and Vicoso 2023; Li et al. 2024). It is currently unclear why turnover has occurred in Diptera but not other insect orders, although one possibility is that the shrinking of gene content on the X that occurred in the ancestor of this group facilitated its reversal to an autosome (Vicoso and Bachtrog 2015; Lasne et al. 2023; Toups and Vicoso 2023). However, turnover was studied using highly fragmented genome assemblies, and a detailed and systematic genomic characterization of derived X chromosomes has yet to be performed. Here, we take advantage of many fly and mosquito chromosome-level genome assemblies that have recently been made available by the Darwin Tree of Life and other projects (Richards et al. 2005; Konganti et al. 2018; Mahajan et al. 2018; Bracewell et al. 2019; Hill et al. 2019; Renschler et al. 2019; Hemmer et al. 2020; Mai et al. 2020; Massey et al. 2020; Small et al. 2020; Torosin et al. 2020; Chakraborty et al. 2021; Durkin et al. 2021; Urban et al. 2021; Zamyatin et al. 2021; Lukyanchikova et al. 2022; Romine et al. 2022; The Darwin Tree of Life Project Consortium 2022; Wei et al. 2022; Alencar et al. 2023; Leung et al. 2023; Meisel et al. 2023; Reinhardt et al. 2023; Tandonnet et al. 2023; Trinca et al. 2023; Congrains et al. 2024) to confirm several putative sex chromosome transitions, as well as identify several additional ones. We also investigate some genomic and evolutionary features of these independently derived X chromosomes that could not be studied previously, such as the fate of reverted X chromosomes and the role of gene movement in shaping the gene content of the X and autosomes.

## Results & discussion

### 1 Additional sex-chromosome turnover events and X-to-autosome fusions

Earlier work identified sex-chromosome turnover (with the reversal of element F to an autosome) using highly-fragmented genome assemblies; scaffolds were assigned to the X or autosomes based on coverage, but their location on different autosomes was unknown (Vicoso and Bachtrog 2015). There are now over 100 dipteran species for which chromosome-level assemblies with an identified X are available. We mapped the coding sequences of *D. melanogaster* to each chromosome of 149 dipteran genomes (listed in Table S1) to identify their homology to Muller elements (see Figure S1 for the graphical representation of these genes in the chromosomes of one species per family). While fewer homologs were detected in species that are more distant from *Drosophila*, more than 6000 were identified in each, giving us sufficient power to identify chromosomal homology. We calculated the expected number of shared genes between each pair of *D. melanogaster* / outgroup chromosomes, and inferred homology when the ratio of the observed to expected number was above 1.5 (following (Toups and Vicoso 2023)), and this excess was significant with a Fisher’s exact test (Table S2). We could then compare the X of different species in a phylogenetic framework (based on the phylogeny of (Sperling and Glover 2023), modified from (Wiegmann et al. 2011)); when several species in a family had the same chromosomal complement, only one was used for further analyses. As expected of an ancestral X, element F corresponded to the sole X in 26 out of 41 families (Figure 1A and Table S1). We recovered several transitions that were inferred previously, including the turnover events of Drosophilidae and Diopsidae (in which the ancestral X reverted to an autosome, e.g. the clades of Figure 1A where element F is no longer part of the X). Interestingly, while the X of *A. gambiae* was previously inferred to derive from a turnover event to element A (Vicoso and Bachtrog 2015), all but two of the *Anopheles* species sampled here (*A. rivulorum* and *A. coluzzii*) show a significant enrichment of both element A and element F genes on the X, suggesting an X to autosome fusion instead (Table S2). A 2-fold enrichment of element F genes is present on the X of *A. coluzzii* (Supplementary Dataset 1), and a large proportion of putative X-linked genes is on unmapped scaffolds in *A. rivulorum* (Supplementary Dataset 2), suggesting that the difference reflects a loss of power in these species rather than biological variation. It should be noted that some lineages, such as Drosophilidae, have been sampled at much higher depth than others, and this may account for the apparent excess of neo-sex chromosome origination events.

**Figure 1:**
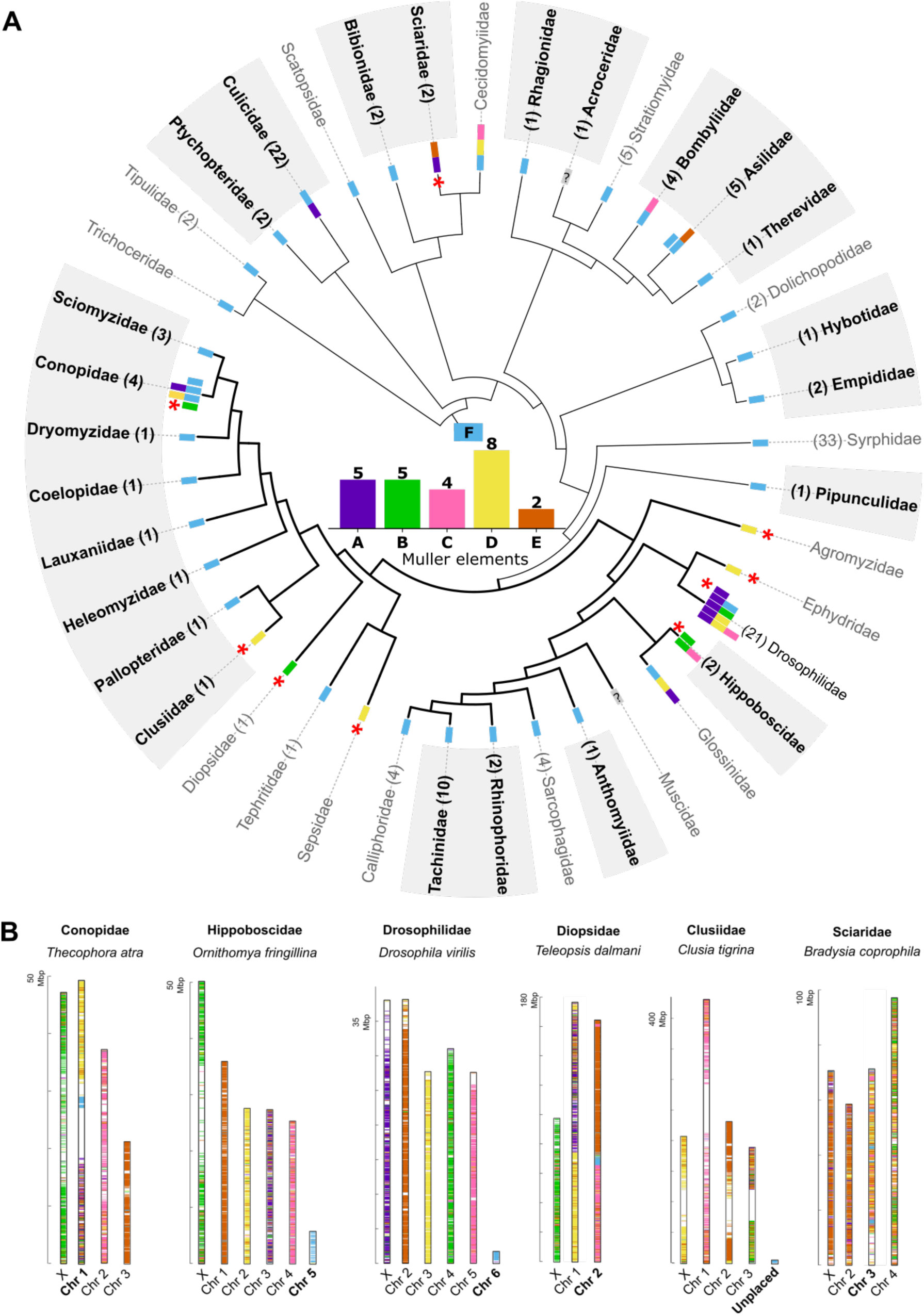
Multiple transitions in sex chromosomes in dipteran insects. A. Phylogeny of the families sampled here, with their corresponding chromosome complement(s). The bars at each edge of the phylogeny represent the Muller elements that are homologous to the X chromosome(s) of that family, either inferred in this study (shaded family name) or previously reported (Vicoso and Bachtrog 2015; Ellison and Bachtrog 2019). Branches highlighted in bold/black correspond to the Schizophora group, and turnover events are marked with red asterisks. The barplot summarizes the number of families in which each Muller element has been co-opted as an X chromosome. Numbers in parentheses next to family names correspond to the number of species analysed in this study. In Polleniidae (which was missing in the reference tree used here, and is therefore not shown) the X is homologous to element F. B. Graphical representation of *D. melanogaster genes* in chromosomes of lineages where a turnover event has occurred (only one species per lineage is represented). Autosomes homologous to F are highlighted.

We also uncovered four additional turnover events in Clusiidae, Conopidae, Hippoboscidae and Sciaridae, and the presence of several additional neo-X arms through fusions to the ancestral X (in Conopidae, Bombyliidae, and Asilidae). It should be noted that although the X chromosome of Sciaridae corresponds to both elements A and E (Figure 1), this represents a simple turnover event, as a homologous autosome containing parts of elements A and E is present in the sister group Bibionidae (chromosome 1 in *Bibio marce*, Figure S2). This is in line with Gries, Jaron and colleagues (*submitted*), who recovered similar sex chromosome transitions when using ancestrally reconstructed chromosomes of Diptera as their reference instead. In total, 9 full turnover events have now been uncovered in this clade, of which 6 happened in groups with chromosome-level assemblies including an annotated X are available (this number does not include turnovers to undifferentiated sex chromosomes (Vicoso and Bachtrog 2015), which we do not consider here). In 3 lineages (Conopidae, Diopsidae and Sciaridae), turnover co-occurred with the fusion of element F to an autosome, while in two (Drosophilidae and Hippoboscidae) element F reverted to an independent autosomal element (Figure 1B and Supplementary Dataset 2). In Clusiidae, element F genes are almost exclusively found in scaffolds that were not assembled into chromosomes (Supplementary Dataset 3), suggesting that a “dot” chromosome may also be present in this lineage. Given that only a small subset of lineages have fusions involving element F (6 out of 30 families where no turnover occurred, Fig 1A), the fact that at least 3 out of 6 lineages that underwent turnover do so suggests that such fusions may favor turnover, or stabilize the existence of the reverted ancestral X after turnover. This is in stark contrast to other clades, where X:autosome fusions consistently lead to neo-X formation rather than turnover (Ellison and Bachtrog 2019; Sigeman et al. 2021; Bracewell et al. 2024; Jayaprasad et al. 2024; Wright et al. 2024).

Turnover events and X to autosome fusions may reflect general dynamics of rearrangements in dipteran insects, or alternatively, may have been favored by selection if they allowed for the sex-linkage of alleles or genes with sex-specific benefits. In particular, sexually-antagonistic alleles, i.e. alleles that benefit one sex at the expense of the other, are theoretically expected to promote the fusion of the autosome they are on to either the X or Y chromosomes (Charlesworth and Charlesworth 1980; Matsumoto and Kitano 2016), and can in principle also favor turnover (van Doorn and Kirkpatrick 2007). If such selective processes are at play, fusions involving sex chromosomes may be more common than expected by chance (Anderson et al. 2020). Clear 1:1 homology between Muller elements is only found in the section Schiphozora (which includes *Drosophila* and other muscoid flies), and therefore we limited our analysis of fusions between X-linked and autosomal elements to this group (Figure S3). We did not consider fusions that occurred in *Drosophila*, as these have been considered elsewhere, and found to be rarer than expected by chance (Anderson et al. 2020). We find only 3 fusions involving the sex chromosome, and 5 autosome-autosome fusions, compared to an expected 2.66 and 5.33 respectively (Blackmon and Adams 2015; Anderson et al. 2020), small numbers that at least do not provide evidence of selection. Similarly, it has been hypothesized that some chromosomes may be more prone to accumulating sexually-antagonistic mutations (if they contain genes that are under strong sexual conflict), and may therefore be co-opted as sex chromosomes more often than expected by chance (Charlesworth and Charlesworth 1980; Marshall Graves and Peichel 2010; Jeffries et al. 2018; Toups et al. 2019; Anderson et al. 2020; Mora et al. 2024). In our dataset (including turnover and autosome-sex chromosome fusions), element D was co-opted as the X in at least 8 families, element A and B in 5, and elements C and E in 4 or fewer families. This distribution is not different from random sampling (p=0.23 of one Muller element being represented at least 8 times with binomial testing). While it is possible that this just reflects a lack of power due to the small number of events, we currently have no evidence of selection favoring one chromosome over the others.

While Diptera lineages generally have highly dynamic sex-chromosome evolution, one family stood out: in 4 species of bee grabbers (Conopidae) for which an annotated X chromosome was available, 4 different X chromosome complements were found (Figure S4). *Conops quadrifasciatus* maintains the ancestral X, while fusions to this chromosome happened in *Myopa testacea* (F+D) and *Sicus ferrugineus* (F+A, although it is likely that this represents a Y-autosome fusion, Layana et al., under preparation). Finally, a turnover event occurred in *Thecophora atra*, where element B replaced the original element F as the X. The time for the divergence of these Conopidae species is only 44 million years, highlighting how dynamic sex chromosomes are in this group.

### 2 The section Schizophora has reduced gene content on element F and increased turnover rates

One hypothesis for why dipterans undergo recurrent sex-chromosome turnover is the shrinking of the X that occurred in their ancestor (from >1000 to ∼100-200 genes) (Vicoso and Bachtrog 2015; Toups and Vicoso 2023). However, even within Diptera, the proportion of *Drosophila* genes that map to the chromosome homologous to the element F can vary by a factor of ∼5 (Table S3), from 0.66% of all mapped genes in Calliphoridae to >3% in Ptychopteridae (Figure 2). It is therefore possible that, even within this order, turnover may be shaped by this difference in gene content. In particular, 8 out of 9 of the turnover events (with reversal of element F to an autosome) happened in the section Schizophora, despite a small difference in the total number of families that were sampled inside and outside of the section (21 vs 18). To test if the rate of turnover is higher in Schizophora than in other dipteran insects, we produced a rooted tree with branch lengths using available gene annotations, as well as BUSCO orthologs for lineages for which an annotation was not available (Table S4). We then applied the approach of (Revell et al. 2024) to compare the log likelihood of models of turnover with homogeneous rates throughout the phylogeny to the log likelihood of equivalent models that allow for variation between Schizophora and other branches (Figure S5A). Both types of models confirm the intuition that high turnover rates are a feature of Schizophora (p=0.004 for “ER” models, which allows for equal rates of turnover and turnover reversal, and p=0.007 for “ARD” models, which assumes different rates of turnover and reversal, a more realistic model in our case; both p-values were obtained with likelihood ratio tests).

**Figure 2:**
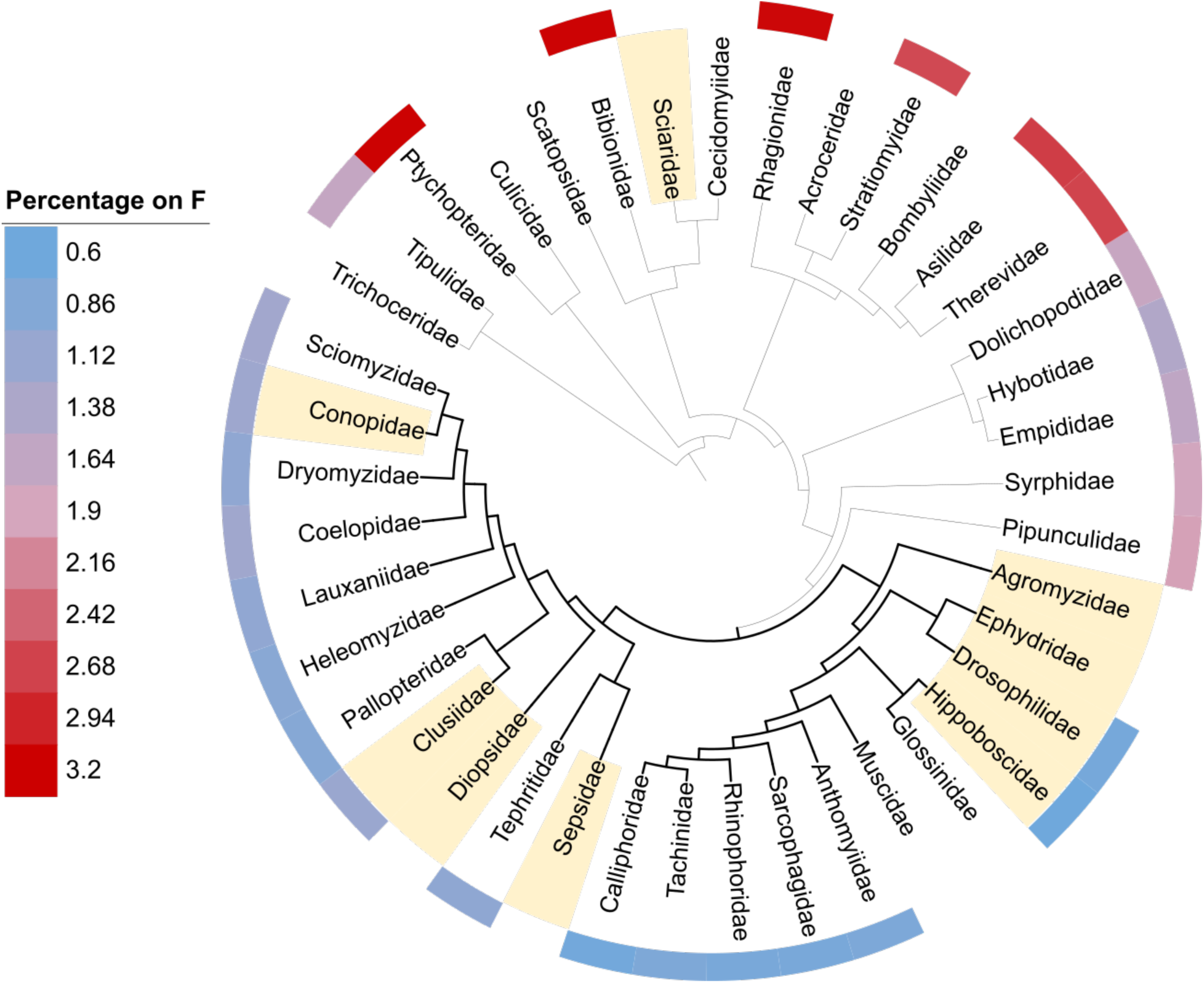
The proportion of genes located on the ancestral X (element F) is variable across Diptera (outer circle). Shaded edge family labels denote sex chromosome turnover (with reversal of the F to an autosome, either fused or unfused to another Muller element). Branches shaded in black correspond to the Schizophora section, where 8 out of 9 turnovers occurred.

All Schizophora families still using element F as their X carry fewer than 1.25% of mapped genes on this chromosome, suggesting that this small proportion may be ancestral to the clade. Outside of Schizophora, on the other hand, between 1.45 and 3.19% of mapped *Drosophila* genes are on element F. We used the Phytools function fastAnc to perform ancestral reconstruction of the percentage of genes on F throughout the phylogeny (Figure S5B and Supplementary Dataset 4). This analysis confirms that shrinkage of the F occurred in the ancestor of Schizophora (1.12% of genes on F, 95% confidence interval: 0.89-1.50%), with all nodes within Schizophora having smaller percentages than those outside of Schizophora (p=7.4e-07). These results therefore support the concurrent shrinking of the gene content of element F and an increase in sex chromosome turnover. While this association may be due to chance or sampling biases, better sampling outside of Schizophora may uncover more fine-grained heterogeneity that could contribute to turnover elsewhere. Finally, while entirely anecdotal, it is intriguing that the smallest chromosome homologous to element F is found in Hippoboscidae (0.6% of mapped genes), where it reverted to an independent dot-like autosome, again in line with the idea that small chromosomes can undergo turnover.

### 3 Muscidae: the mysterious case of the disappearing element F

In most dipteran lineages, element F was conserved as the sex chromosome, reverted to an autosome, or was incorporated into an autosome through a fusion. In *Polietes domitor* (Muscidae), on the other hand, no chromosome showed strong evidence of homology to element F (Figure 3C). The putative X chromosome of this species carries too few genes (1) to determine homology to a Muller element. To investigate what might have happened to element F in this clade, we first searched for element F genes in the 10 species of the family Muscidae for which genomes were available, even if they did not have an assigned X chromosome in the assembly. In two species of the genus *Hydrotaea* (Falk and Grzywacz 2024; Falk and Grzywacz 2024), small scaffolds (assigned as B1 and B2 in *H. diabolus*, and as chromosome 6 in *H. cyrtoneurina*) were enriched for element F genes (Figure 3B and Figure S1). However, these scaffolds were much smaller than chromosomes homologous to element F in other families, with a total of only 17 and 20 genes mapped in the two species (of which 7 were common to both).

**Figure 3:**
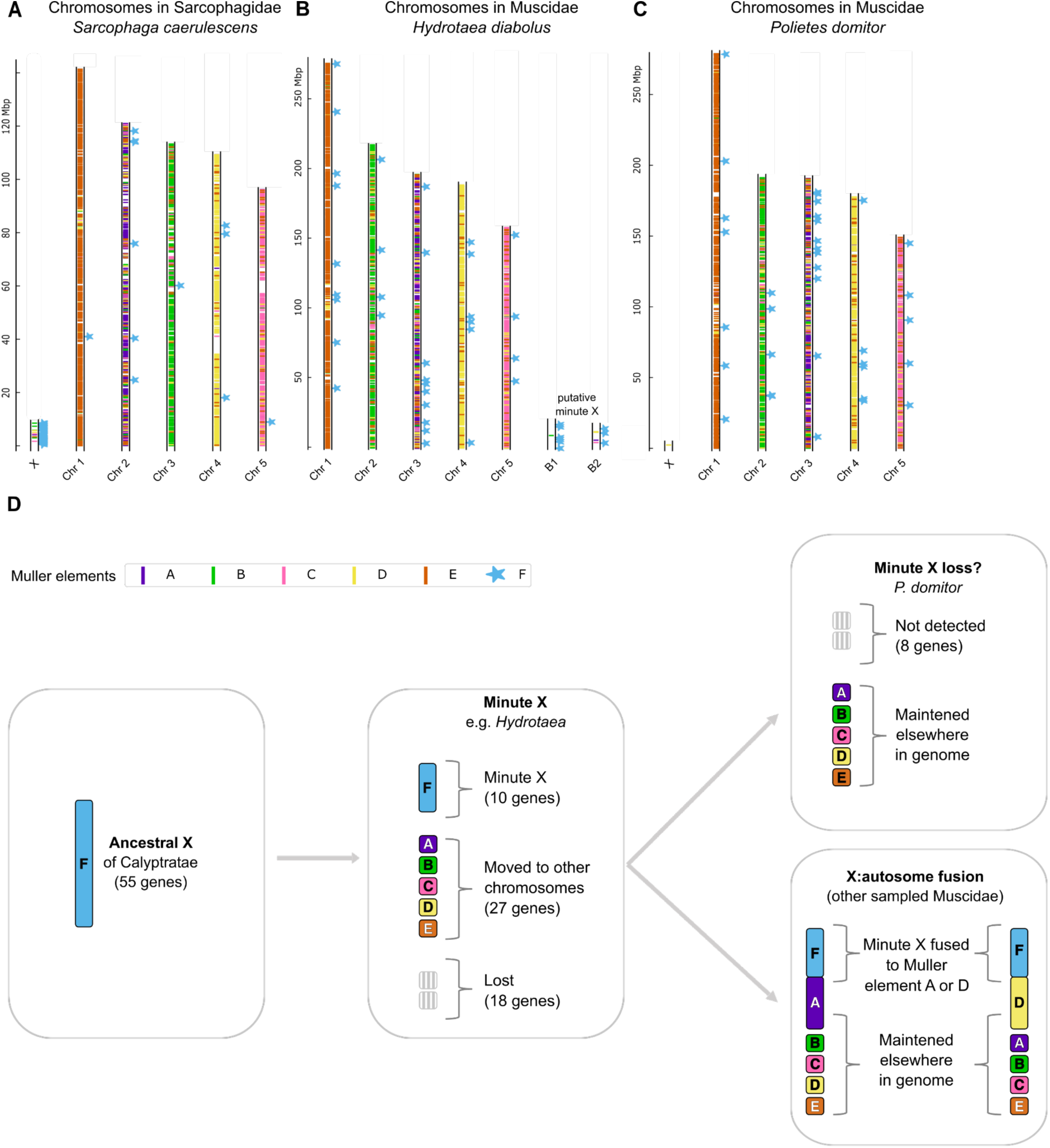
Extreme shrinking of Muller element F in Muscidae. **A.** Genomic location of *D. melanogaster* genes mapped to the genomes of *Sarcophaga caerulescens* (Sarcophagidae), colored according to their Muller element in *D. melanogaster,* to show an example of a chromosome homologous to element F (the X chromosome); **B.** the same for *Hydrotaea diabolus* and **C.** for *Polietes domitor* (Muscidae). **D.** Model for the evolution of Muller element F in Muscidae.

Karyotypic data in other *Hydrotaea* suggests that in addition to having 5 large euchromatic chromosomes, species are often polymorphic for small heterochromatic chromosomes, with their number varying from 1 to 7 across individuals and species (Loeschcke et al. 1994). The two small scaffolds of *H. diabolus* had reduced coverage in the sequenced male individual, and were in light of this karyotypic data inferred to be supernumerary (B) chromosomes (Falk and Grzywacz 2024). However, such a coverage pattern is equally likely to reflect a small X chromosome (or two small X chromosomes), a simpler explanation to account for both reduced coverage and homology to the ancestral X. It is further possible that such a small X could vary in number across cells, as a similar pattern has been found for the dot chromosome in *Drosophila melanogaster* (Mohr 1932). We therefore hypothesize that the ancestral X underwent dramatic shrinking in the ancestor of Muscidae to form the minute X of *Hydrotaea*. In other Muscidae species, no such small chromosomes with homology to element F were reported.

To further investigate the evolution of element F in Muscidae, we selected genes that are on F in *D. melanogaster* and on its homologous X in *Sarcophaga caerulescens* (Figure 3A), and which therefore must have been on the ancestral X of the lineage leading up to Muscidae (Calyptratae) (Table S5). Of these 55 genes, only 10 (18%) were found on the minute X chromosomes of either *Hydrotaea* species. 18 genes (32%) cannot be found in any of the 10 Muscidae species, suggesting that the shrinking of Muller element F in the ancestor of this group was accompanied by substantial gene loss. A similar pattern is observed when all *D. melanogaster* genes from element F are mapped to each of the Muscidae species, with a relatively large proportion missing (Figure S6: 49-62% of element F genes recovered, compared to >80% in other comparably distant lineages). Finally, 27 of the 55 ancestrally X-linked genes (49%) are detected in at least one Muscidae species, but are not found on the *Hydrotaea* minute X, suggesting that extensive gene movement from the X to the autosomes also contributed to shrinking element F.

The 10 minute X genes had different fates in the various Muscidae that did not have an independent Muller element F (Figure 3D). In *Eudasyphora cyanicolor* and *Muscina levida*, the majority of them were found on Muller element A (8/8 and 6/9, respectively), suggesting that a minute X - A fusion occurred at least once. In the stable fly *Stomoxys calcitrans*, all 8 out of 8 minute X genes were instead found on Muller element D, supporting an independent fusion of the minute X to this chromosome. These observations are in line with (Meisel et al. 2020), who inferred that the X chromosomes of the horn fly and the stable fly correspond to A-F and D-F fusions respectively. Interestingly, this work (which was based on partly assembled genomes) already noted the small number of genes putatively assigned to element F, as well as the lack of an observed homologue of element F in cytological studies. Finally, in *Polietes domitor,* only 2 out of 10 minute X genes were recovered (one on element D, and one on an unmapped scaffold). While it is possible that a minute F is found in this species but failed to assemble, it is also possible that this small chromosome was lost in this lineage (the BUSCO score for the assembly is >99%, (Falk et al. 2024)). Muscidae lineages are very labile in their sex chromosome complements (Meisel et al. 2020). The house fly *Musca domestica* is also noted for its highly polymorphic sex chromosomes, with all of the 5 large chromosomes having been found to carry the male-determining factor (Sharma et al. 2017). Muscidae may therefore perfectly exemplify how an extreme reduction in gene content of the X can be accompanied by highly variable sex determination systems, although more work to identify the X chromosome in its various lineages is still needed to fully characterize turnover dynamics in this clade.

### 4. High GC content on derived X chromosomes supports widespread male achiasmy

Most *Drosophila* species are male-achiasmatic, i.e., meiotic recombination typically does not occur in males. Cytological observations in a few other dipteran species suggests that a lack of crossovers in male meiosis may be widespread in this group (Satomura et al. 2019). Male achiasmy dramatically influences sex chromosome evolution, as full sex linkage of a chromosome is expected to initially favor turnover (by linking male-favorable mutations to males), but should also promote rapid degeneration of whole Y chromosomes once turnover has occurred (Satomura et al. 2019). Since measuring recombination rates in non-model species remains challenging, how widespread male achiasmy is remains an open question, which we address here indirectly. In particular, the GC content throughout the genome is typically correlated with the local recombination rate (Charlesworth et al. 2020). In species where recombination is similar in males and females, the X has lower recombination rates than the autosomes because it does not recombine in males (where it is paired with the Y). However, in male-achiasmatic species, the X is expected to have higher rates of recombination than the autosomes, because it spends less time in nonrecombining males. We therefore predict that if male achiasmy is widespread, novel X chromosomes should overall have acquired a higher GC content than autosomes.

We compared the GC content of the X and autosomes of the 11 lineages that acquired one of the large Muller elements as the X (either by itself or as a fusion to the ancestral X, as the small size of element F means the signal will likely be driven by the neo-X chromosome). Specifically, chromosomes were broken into 100kb windows, which were compared. It should be noted subdividing chromosomes into windows could introduce pseudoreplication, and the absolute p-values should therefore be interpreted with caution. For lineages for which several assemblies were available, only one is represented in Figure 4 (see Supplementary Dataset 5 for GC content estimation in all the species). In 7 out of 11 lineages, the X had significantly higher GC content than the autosomes, consistent with lack of recombination in males being common in Diptera. In *D. melanogaster* and in *Sicus ferrugineus* (Conopidae) the GC content of the X was similar to that of the autosomes, while in only two species it had significantly lower GC content (Sciaridae: *Bradysia coprophila*, Bombyliidae: *Bombylius discolor*). At least one Sciaridae species appears to be male-achiasmatic (Amabis et al. 1979). However, Sciaridae males do not transmit the paternal copy of their genome (neither the X nor the autosomes) (Baird et al. 2025), and sex-averaged rates of recombination should therefore be equal for all chromosomes in this group. Bombyliidae, on the other hand, may represent a true case of loss of male heterochiasmy, as low GC content on the X is consistent in the 4 available species. The fact that *D. melanogaster*, the best known example of male achiasmy, does not have higher GC content on the X also shows that other factors play a role (such as differences in female recombination rates between the X and autosomes, or sex-biased mutation rates). This is in line with previous data that recombination rate and GC content are negatively correlated on the X in *D. melanogaster*, though why this occurs is not fully understood (Singh et al. 2005). Despite these limitations, our data is therefore at least consistent with a prevalent role of male achiasmy in shaping dipteran sex chromosome evolution.

**Figure 4:**
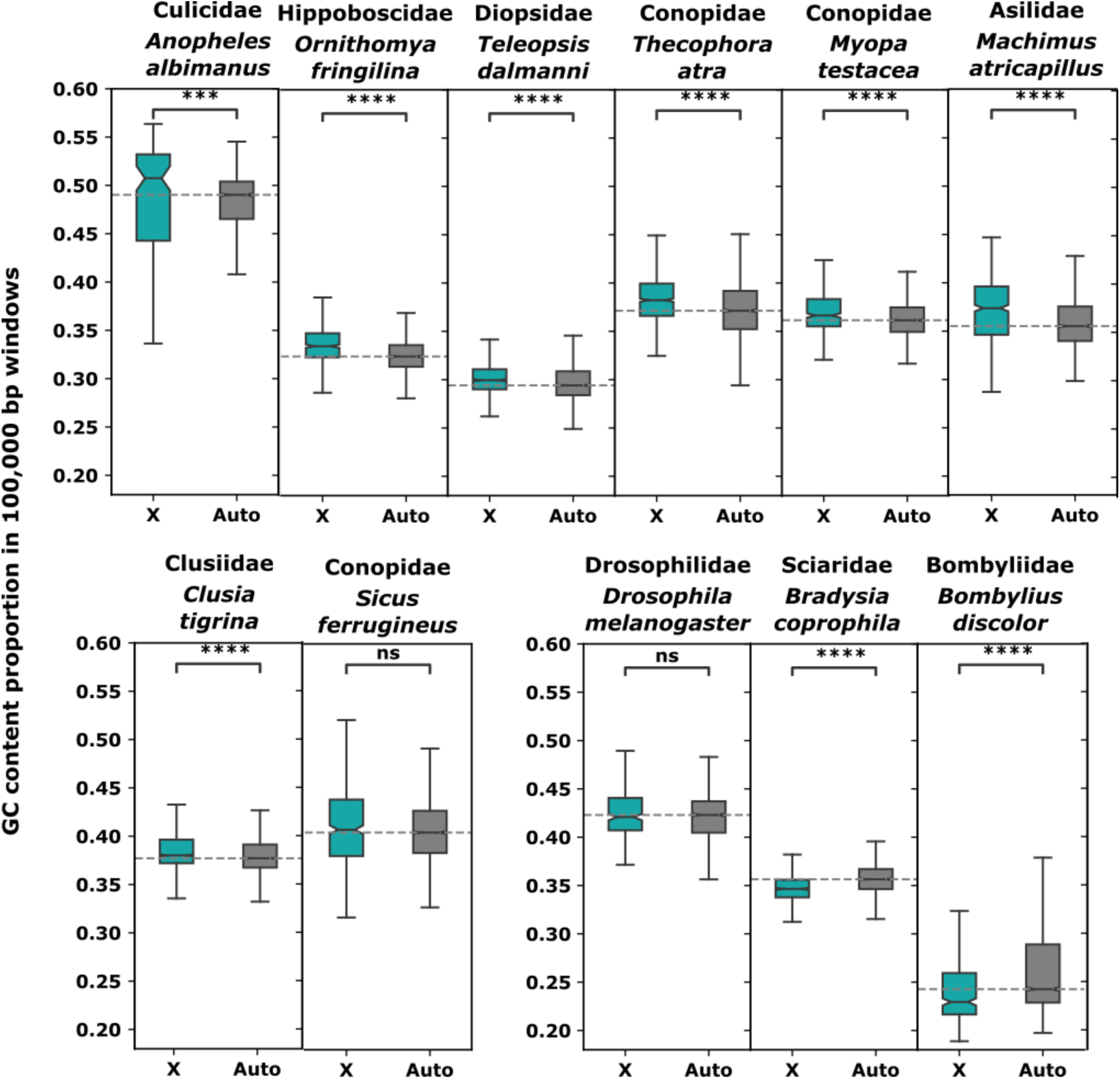
GC content proportion in 100,000 bp windows of the X and autosomes in Diptera species representative of lineages with derived large X chromosomes (p-value annotation legend according to a Mann-Whitney statistical test: **** for p <= 0.0001, *** for p <= 0.001, ** for p <= 0.01, * for p <= 0.05, ns for not significant).

### 5. Excessive movement out of most but not all X chromosomes

X chromosomes often have an unusual gene content. In many insects, they are enriched for female-biased genes, but have a deficit of male-biased genes (Sturgill et al. 2007). How this evolves is unclear, but one potential contributor is the excessive gene movement from the X to the autosomes that has been observed in various clades (Betrán et al. 2002; Emerson et al. 2004; Baker and Wilkinson 2010; Toups and Hahn 2010; Miller et al. 2022). Such runaway genes often have testis- or male-biased expression (Potrzebowski et al. 2008; Vibranovski et al. 2009), and their movement could therefore contribute to the demasculinization of the X (Sturgill et al. 2007). Most of these analyses were done within clades that share the same X chromosome, and relied on duplicates to infer movement, such that the original copy of the genes still remained on the X. We therefore took advantage of the many X chromosomes of dipterans to extend these analyses to see if an excess of genes fully move from the X to the autosomes over longer evolutionary times.

We selected from the phylogeny trios of species where (1) a focal species has a newly derived X that corresponds to one of Muller elements A to E; and (2) two independent outgroups carry element F as their X. To avoid biases due to different qualities of annotations, we mapped the *D. melanogaster* gene sequences and used their best mapping location in each of the other genomes as a proxy for the location of their ortholog. We used the two outgroups to infer the ancestral location of the genes (if it was found on homologous chromosomes in the two, otherwise an ancestral location could not be inferred), and then inferred gene movement if in the focal species the gene was on a different chromosome (see Table S6 for the number of gene movements between chromosomes in each of the trios). Since the correspondence of Muller elements and chromosomes becomes more difficult to assess with increasing distance to *Drosophila*, we focused exclusively on new X chromosomes found in the section Schizophora (in Diopsidae, Clusiidae, Hippoboscidae, Conopidae and Drosophilidae). We selected *D. virilis* as representative species of the Drosophilids because each chromosome comprises a single Muller element, making the analysis more straightforward. In 4 out of 6 species, a clear out-of-X pattern could be seen (Figure 5A), with the X having lost a greater fraction of genes than the other large elements even after controlling for chromosome size, which should affect the target space of gene movement. In the other two, however, the X did not stand out from other chromosomes. While this could reflect true biological variability, it is also possible that these patterns are influenced by the age of these sex chromosomes, as gene loss is likely to be slow. In particular, the strongest effect is seen in *Drosophila*, whose X chromosome has existed for >60 millions of years (estimate of divergence time between *Drosophila* and *Phortica*, with whom the X is shared). On the other hand, all new Conopidae X chromosomes arose since the split of the genera *Conops* and *Myopa*, estimated at 44 million years ago (Kumar et al. 2017), and no estimated age is available for Hippoboscidae. In the two lineages where element F reverted to an autosome without fusing (Hippoboscidae and Drosophilidae), we detect a very strong out-of-F effect, likely due to its history as an X during part of the history of the focal lineages.

**Figure 5:**
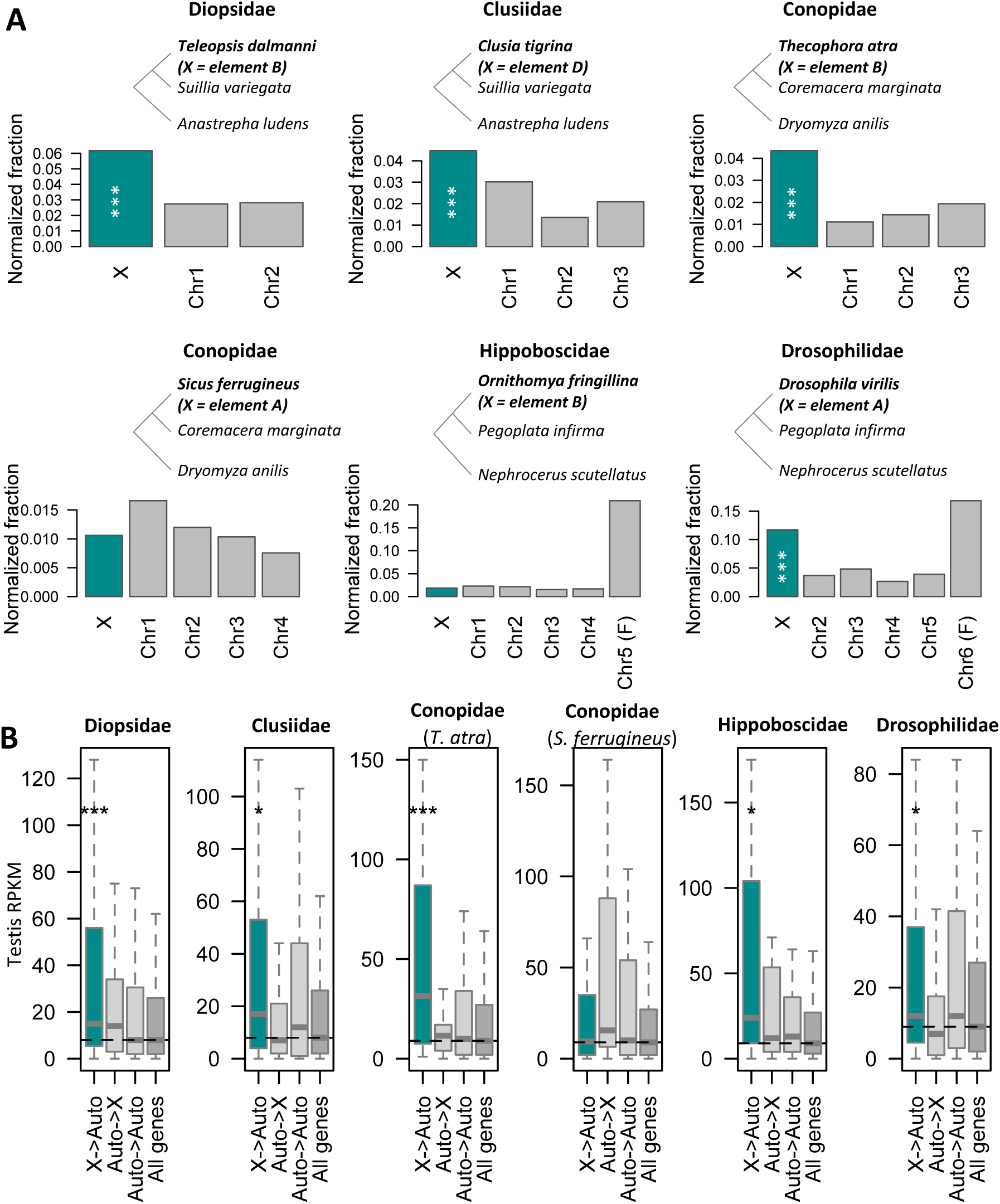
Out-of-X gene movement in Schizophora species which have co-opted a large Muller element as their X chromosome after turnover. **A.** Bar plots show the fraction of genes inferred to have moved out of each chromosome in the focal species, using the trios shown in the phylogenetic trees above each (focal species are in bold). Since smaller chromosomes have a higher chance of experiencing gene loss (because their target space on other chromosomes is larger), we normalize the fractions by dividing them by the fraction of genes found on other chromosomes. The bars representing the X chromosome are in green. Asterisks denote a significant excess of movement out of the X chromosome (*** for p<0.001, ** for p<0.01, * for p<0.05) without correcting for multiple comparisons. All p-values remain significant after Bonferroni correction. **B. *D. melanogaster* testis expression of homologs inferred to have moved chromosomes, and that of all mapped genes, for each of the trios (named after the focal lineage under study)**. X->Auto refers to genes that moved out of the X, Auto->X refers to genes that moved into the X, Auto->Auto refers to genes that moved between autosomes, and All genes refers to all mapped genes in each trio. Significance was assessed with Wilcoxon tests (* for p<0.05, ** for p<0.01, *** for p<0.001, no asterisk for non significant).

We investigated whether genes that moved out of the X had unusual patterns of expression in *D. melanogaster* (as a proxy for ancestral expression, with the exception of the Drosophilidae analysis, where it largely represents current Drosophilid expression). For each of the tissues in the FlyAtlas2 dataset, we compared the distribution of expression of genes that moved out of the X to the expression distribution of all mapped genes. Genes that moved out of the X had significantly higher expression in between 0 and 12 tissues (Wilcoxon tests). In 5 of the 6 trios, genes that moved out of the X had significantly higher expression in testis than all genes (Figure 5B; the only species where this was not found, *Sicus ferrugineus*, also did not show an out-of-X effect), in line with the idea that out-of-X gene movement may deplete X chromosomes of genes with male-biased expression. This was not the case for genes that moved into the X, or between autosomes (p>0.05 in all, Figure 5B). Ovary expression was also enriched in out-of-X genes in 3 lineages (Figure S7), but in two of those (Drosophilidae and Conopidae) less so than testis expression. The one exception is Hippoboscidae, where out-of-X genes had much stronger ovary expression than any other category of genes; this lineage also did not show an out-of-X effect. Both testis and ovary enrichment patterns held when modENCODE expression values were used instead (Figures S8-S10), except for the high ovary expression of out-of-X genes in Drosophilidae, which was not detected in these data. Finally, we plotted expression distributions for somatic controls (female and male head, female and male carcass, Figure S11-S14), where out-of-X genes were never overexpressed compared to the full set of mapped genes. Taken together, these results suggest that the high expression observed for gonads does not simply represent broadly high expression, and that in lineages where an excess of movement from the X is observed, these moved genes tend to have high testis expression.

We also performed a functional enrichment of genes that had moved out of the X using stringDB with the *D. melanogaster* genome as background (Szklarczyk et al. 2023). In the two lineages where no excess of gene movement could be detected, no enrichment was found either. In the others, between 11 and 40 functional terms were enriched (FDR<0.05, see Table S7-S11), again suggesting that the set of genes that move out of the X is not random. Interestingly, several of these functional terms were shared by at least two species (Figure S15). When the genes that were mapped in each trio were used as the respective background instead (a more conservative approach to control for biases in what genes are mapped across species), 4 terms related to metabolism were enriched in Drosophilidae (Table S10), and one in Diopsidae (RNA polymerase, Table S7). This reduced number of functional enrichments may reflect a loss of power due to the smaller background set, but also suggests some caution in interpreting the larger sets of functional enrichments. Taken together, these results show that the out-of-X pattern is not unique to the lineages previously sampled, and that the specialized set of genes that move is likely to shape X chromosome gene content over long evolutionary time periods.

## Conclusions

We analyzed more than 100 different chromosome-level genome assemblies of Dipterans, and identified 4 new sex-chromosome transitions from the ancestral insect X in Diptera (in addition to the 5 previously identified). Similar to previous analyses of *Drosophila* sex chromosomes, transitions to new sex-chromosome systems did not preferentially involve a single Muller element, suggesting there was no single element predisposed to sex-linkage. Interestingly, most turnovers occurred within the Schizophora, which also has fewer genes on Muller element F than lower Dipterans, suggesting that even on this scale reverting the ancestral X chromosome to an autosome may be less deleterious with fewer genes. We indirectly tested for male achiasmy in the species with novel X chromosomes. We found that most of these X have higher GC content than the autosomes, supporting widespread male achiasmy in Diptera. Finally, excess gene movement off the X is common and, over time, may contribute to the depletion of testis-expressed genes that is typically observed on old X chromosomes.

## Methods

### Genome assemblies

We downloaded from NCBI (https://www.ncbi.nlm.nih.gov/assembly) the set of available Diptera genome assemblies, and their respective list of chromosomes. We selected all species for which “X” was found in a chromosome description, and manually checked that an X chromosome had been annotated. While this search was first performed in February 2023, further species were added to the dataset as they became available. The full list of analysed genomes, as well as their respective publication when available, is provided in Table S1 (including scaffolded genomes and genomes with no annotated X in Muscidae and Conopidae).

### Phylogenetic tree

A phylogenetic tree of Diptera comprising all but one of the families for which a genome was analysed was adapted from (Sperling and Glover 2023), and visualized with iTOL, version 7.1 (Letunic and Bork 2024). iTOL was also used to edit the phylogenetic trees shown in Figure 2A and Figure S3 and S4. The published phylogeny did not include branch lengths. In order to be able to perform the phylogenetic inferences described in section “**2 The section Schizophora has reduced gene content on element F and increased turnover rates**”, we downloaded protein annotations for one representative of each family from Ensembl (Darwin Tree of Life) https://projects.ensembl.org/darwin-tree-of-life/ or other sources (Table S4) and extract the longest sequence per protein using a custom Perl script (GetLongestCDS_V2.pl). When no annotations were available (in the case of *Glossina fuscipes* (Glossinidae), *Liriomyza trifolii* (Agromyzidae), *Cirrula hians* (Ephydridae species), *Sitodiplosis mosellanafa*, (Cecidomyiidae), *Coboldia fuscipes*, (Scatopsidae), Trichoceridae species, *Coelopa pilipes* (Coelopidae), *Dryomyza anilis* (Dryomyzidae), *Ornithomya fringillina* (Hippoboscidae) and *Rhagio lineola* (Rhagionidae), we ran BUSCO version v5.8.2 (Manni et al. 2021) on the genome assembly with the Diptera gene set, and used the resulting conserved genes instead (we used the default gene mapper Miniprot (Li 2023) but for *C. hians*, we used Metaeuk (Levy Karin et al. 2020) as the gene predictor. The predicted genes were then passed to HMMER (Eddy 2011)). In the case of *C. hians*, we used an improved assembly that was obtained by running Megahit (Li et al. 2015) with default parameters on all available published Illumina genomic reads (NCBI Bioproject accession: PRJNA268392). We also included the outgroup *Panorpa germanica* (protein set from Ensembl, see Table S4). All protein sets were used as input for Orthofinder (with standard parameters), which produced a rooted species tree.

### Mapping of *Drosophila* genes to outgroup genomes

*D. melanogaster* genes were retrieved from Ensembl (https://nov2020.archive.ensembl.org/Drosophila_melanogaster/Info/Index). We filtered for the longest sequence of each CDS using a custom Perl script (GetLongestCDS_V2.pl). We then used Standalone BLAT, version 36x2 (Kent 2002) to map the resulting gene sequences to the dipteran genomes using default parameters but with translated DNA for the query and database. We kept the best hit for each gene and kept only those genes that do not overlap or overlap by <20 bp using two Perl scripts (1-besthitblat.pl and 2-redremov_blat_V2.pl). *D. melanogaster* genes were mapped to all the dipteran genomes specified in Table S1. The set of *D. melanogaster* genes and their assigned chromosomal location in each outgroup genome is provided as Supplementary Dataset 7, and the graphical representation of the location of *D. melanogaster* genes colored according to their original Muller element is provided in Supplementary Dataset 2.

### Detection of X-linked Muller elements in dipterans

We used the results from Blat (described above) to infer X-linked Muller elements in each species. In the absence of homology, the expected number of genes that should be shared by a pair of chromosomes of two species is:

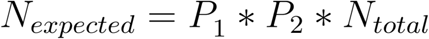

Where P1 is the proportion of genes found on the chromosome of species 1, P2 is the proportion of genes found on the chromosome of species 2, and N_total is the total number of genes found in both species. Chromosomes of outgroup species were inferred to be homologous to *Drosophila* Muller elements when the number of genes they shared with the Muller element was greater than 1.5-fold the expected number.

### Detection of X-autosome and autosome-autosome fusions

Within the Schizophora, each Muller element typically corresponds to only 1 chromosome (one exception is found in *Sicus ferrugineus*, where element C is found on two chromosomes; since it is difficult to separate a fission from a misassembly, we consider the two to be a single chromosome in our analyses). On the other hand, Diptera chromosomes can be enriched (>1.5x) for more than 1 Muller element, showing that fusions have occurred. Most lineages only had independent fusions that could be counted directly. Where fusions were present in more than one species in a single family, parsimony was used to infer the number of times they arose.

### Conopidae tree

The Conopidae phylogenetic tree was inferred with Orthofinder, version 2.5.5 (Emms and Kelly 2019), using the multiple sequence alignment option (MSA). This option infers a maximum likelihood tree, which uses MAFFT, version 7.505 (Katoh et al. 2002) to align the sequences and FastTree (Price et al. 2010) to infer the tree. We used protein sequences from the six Conopidae species and a Lauxaniidae (*Calliopum simillimum*) which was used as the outgroup. The fasta files were downloaded from https://projects.ensembl.org/darwin-tree-of-life/ and we filtered for the longest sequence of each protein with the Perl script GetLongestCDS_V2.pl.

### Phylogenetic analyses

#### Comparison of turnover rates within and outside Schizophora

The Phytools (Revell 2012) functions fitMk and fitmultiMk were used to compare the log-likelihood of models that assume a single turnover rate for the whole tree (fitMk) to models that assume two different turnover rates inside and outside of Schizophora (fitmultiMk). Two types of models were used: ER assumes turnover and turnover reversal happen at equal rates, while ARD allows for asymmetric turnover and reversal rates. P-values were obtained with likelihood ratio tests.

#### Ancestral reconstruction of % of mapped genes on element F

For each species in our analysis, we detected the chromosome that was enriched for element F genes (>1.5 fold relative to the expected number, see description of the approach for detecting homology to X). If this chromosome was also enriched for genes from another Muller element, a fusion was inferred, and the species was excluded from this analysis. For each species where no fusion was found, we extracted the total number of genes mapped to the chromosome homologous to F, and divided it by the total number of mapped genes to obtain a fraction of genes on F (Supplementary Dataset 4). We then used the Phytools function FastAnc to perform ancestral reconstruction of the percentage of mapped genes located on element F throughout the dipteran phylogeny. Families for which only scaffold level assemblies were available, or where element F was fused to one of the large chromosomes, were excluded from this analysis (as it was not possible to infer the % of genes on F).

### Estimation of GC content

GC content was estimated in the same way described in (Lasne et al. 2023). In short, for the GC content we used the Python script GCcalc.py (https://github.com/WenchaoLin/GCcalc) in 100,000 bp window size and 100,000 bp step size along the genome of each species.

### Gene movement inference and statistical testing

We focused on the section Schizophora to examine movement of genes between chromosomes, as within this section homologous chromosomes are generally well conserved, making inferences of gene movement reliable. For each species that coopted one of the large Muller elements as an X chromosome, we manually selected two independent outgroups that still had element F as the ancestral X. We first identified homologous chromosomes between the outgroup and focal species. To do so, we compared the mapping location of *D. melanogaster* genes on the chromosomes of each focal-outgroup pair of species, and classified two chromosomes as homologous when they had a more than 1.5-fold excess of observed / expected shared genes (where the expectation is simply: proportion of mapped genes on the chromosome of species 1 * proportion of mapped genes on the chromosome of species 2 * total number of mapped genes). When the location was not in agreement in the two outgroup species, the gene was not considered further; when it was, this location was assigned as its putative ancestral location. Gene movement was then inferred when the location in the focal species was different from the inferred ancestral one.

In order to determine if there was a significant excess of movement out of the X for each species, we computed the expected number of movements between chromosomes using the following formula (taken from (Toups et al. 2011), originally proposed by (Betrán et al. 2002)):

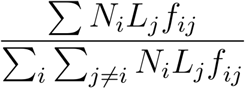

This formula accounts for the number of genes on the chromosome of the parent copy, the size of the chromosome with the daughter copy, and whether movement occurred to an X chromosome or an autosome. The indices i and j are for the chromosomes with the parent copy and with the daughter copy, respectively, Ni is the number of genes on the chromosome with the parent copy, Lj is the length of the chromosome of the daughter copy, and fij is 1 when j is an autosome and 0.75 when j is an X chromosome. We computed the expected gene movements for each chromosome, and then binned the movements as (1) autosome to autosome (2) autosome to X and (3) X to autosome. We then determined significance using a Chi-square goodness of fit test.

### Gene expression analysis

A table containing all the expression values stored in FlyBase (Larkin et al. 2020) was downloaded from their website (https://s3ftp.flybase.org/releases/FB2025_04/precomputed_files/genes/gene_rpkm_matrix_fb_2025_04.tsv.gz) and subsetted for columns containing FlyAtlas2 expression (Leader et al. 2018) or for modENCODE expression (modENCODE Consortium et al. 2010). For each FlyAtlas2 tissue, the expression of *D. melanogaster* homologs of genes that moved out of the X in each trio was compared to the expression of all mapped tissues with a Wilcoxon test (the resulting p-values are provided in Supplementary Dataset 6). For tissues of interest (testis, ovary, male and female head, male and female carcass), comparisons were also performed between genes that moved into the X and the full mapped set, and between genes that moved between autosomes and the full mapped set.

### Functional enrichment on StringDB

The Flybase accessions of *D. melanogaster* homologs of genes that moved out of the X in each of the focal species (previous section) were used as input for StringDB (Szklarczyk et al. 2023) with multiple proteins. Either “*Drosophila melanogaster*” was selected as the reference species, or the set of D. melanogaster genes that were mapped in each of the trios was given as the reference background instead. The enriched functional categories obtained with the standard StringDB analysis with default parameters (as of May 2025) were then downloaded (Table S7 to S10). A similar analysis was performed with sets of genes inferred to have moved between autosomes (Table S11 and 12 for Hippoboscidae and Drosophilidae; no enrichment was found in other lineages).

## Data availability

Supplementary datasets and Table S1, S2, S5 and S6 are available at: https://seafile.ist.ac.at/d/6e69e6be57374552acfe/ [*a permanent URL will be provided upon acceptance*]. Pipelines are available at https://git.ista.ac.at/llayanaf/transitions_diptera.

## Author contributions

MAT and BV designed the original study. LL performed most analyses with support from MAT and BV. LL, MAT and BV wrote the manuscript.

## Funding

This work was supported by a grant from the Austrian Science Fund (FWF, grant number PAT 8748323) to BV.

## Conflict of interest statement

The authors declare no conflict of interest.

## Acknowledgements

We thank the Vicoso group for their feedback on an early version of the manuscript. We are grateful to Kamil Jaron and Julia Gries for helpful discussions and for sharing their unpublished work. Computational resources and support were provided by the Scientific Computing Unit at ISTA.

## Notes

https://seafile.ist.ac.at/d/036324e37c474706a91a/

### Competing Interest Statement

The authors have declared no competing interest.

### Summary of Updates

Revisions made as required by reviewers.

