## Supplement for "Causes and consequences of sex-chromosome turnovers in Diptera"

**Supplementary Materials**

**Supplementary Figures**

**Figure S1:** Graphical representation of *D. melanogaster genes* in chromosomes (only one species per family is represented).
**Figure S2:** Heatmap of Muller elements enrichment in *Bradysia coprophila* and enrichment of *B. coprophila* chromosomes in *Bibio marce*.
**Figure S3** : Fusions in the section Schizophora.
**Figure S4:** The X chromosome composition of Conopidae species and fusions.
**Figure S5:** Schizophora have increased turnover rates and a smaller proportion of genes on element F.
**Figure S6:** Fewer element F genes are recovered in Muscidae genomes (circles) than in other dipteran genomes (triangles) once divergence has been accounted for.
**Figure S7:** *D. melanogaster* ovary expression of homologs inferred to have moved chromosomes.
**Figure S8:** *D. melanogaster* testis expression of homologs inferred to have moved chromosomes, using ModEncode data obtained from mated 4 day old males.
**Figure S9:** *D. melanogaster* ovary expression of homologs inferred to have moved chromosomes, using ModEncode data obtained from mated 4 day old females.
**Figure S10:** *D. melanogaster* ovary expression of homologs inferred to have moved chromosomes, using ModEncode data obtained from virgin 4 day old females.
**Figure S11:** *D. melanogaster* female carcass expression of homologs inferred to have moved chromosomes.
**Figure S12:** *D. melanogaster* male carcass expression of homologs inferred to have moved chromosomes.
**Figure S13:** *D. melanogaster* female head expression of homologs inferred to have moved chromosomes.
**Figure S14:** *D. melanogaster* male head expression of homologs inferred to have moved chromosomes.
**Figure S15:** StringDB functional enrichment of genes that moved out of the X.

**Supplementary Tables** (https://seafile.ist.ac.at/d/6e69e6be57374552acfe/; a permanent URL will be provided upon publication)

**Table S1**: List of analysed genomes and corresponding citations.
**Table S2**: Enrichment of ME acting as the X.
**Table S3:** The proportion of mapped genes found on the chromosome homologous to element F.
**Table S4:** Protein annotations used for our Diptera-branched tree.
**Table S5**: The fate of genes ancestrally located on the X of Calyptratae.
**Table S6**: Number of gene movements.
**Table S7:** GO enrichment amount genes that moved out of the X in Diopsidae.
**Table S8:** GO enrichment amount genes that moved out of the X in Clusiidae.

**Table S9:** GO enrichment amount genes that moved out of the X in Conopidae (*Thecophora atra*).
**Table S10:** GO enrichment amount genes that moved out of the X in Drosophilidae.
**Table S11:** GO enrichment amount genes that moved between autosomes in Hippoboscidae.
**Table S12:** GO enrichment amount genes that moved between autosomes in Drosophilidae.

**Supplementary Datasets** (permanent URL will be provided upon publication)

**Supplementary Dataset 1:** Heatmaps for enrichment of Muller elements in outgroups.
**Supplementary Dataset 2:** Graphical representation of the location of *Drosophila melanogaster* genes colored according to their original Muller element in each outgroup genome.
**Supplementary Dataset 3:** Gene counts. Number of *D. melanogaster* genes assigned to each chromosome in outgroup species.
**Supplementary Dataset 4:** Number and proportion of genes mapped to chromosomes homologous to element F.
**Supplementary Dataset 5:** GC content of all species with an X.
**Supplementary Dataset 6:** Out-of-X-FlyAtlas2
**Supplementary Dataset 7:** *Drosophila melanogaster* genes and their assigned chromosomal location in each outgroup genome.

### Supplementary Figures

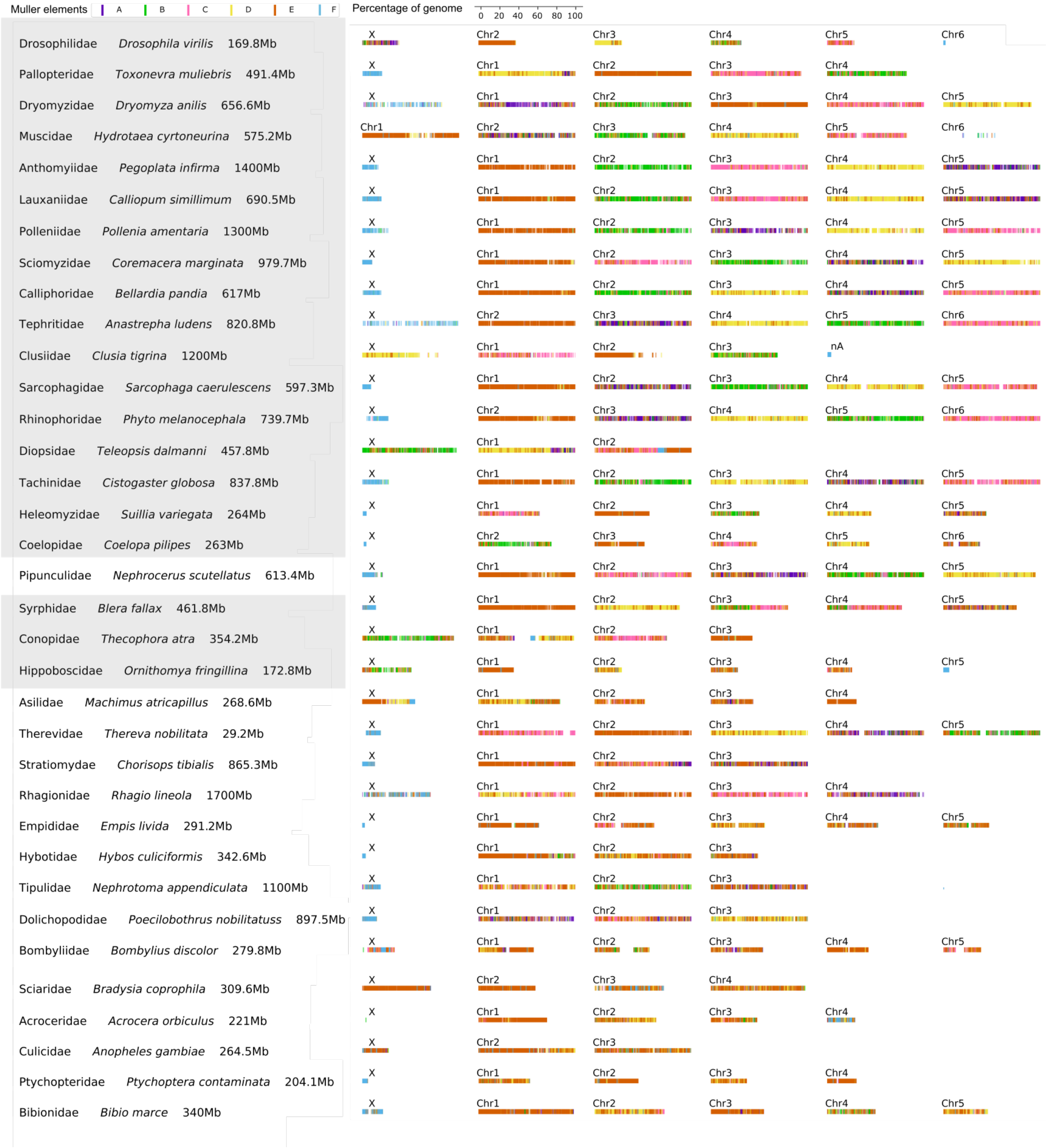

**Figure S1:** Graphical representation of *D. melanogaster genes* in chromosomes (only one species per family is represented). Each chromosome in the species has been normalized by the size of the corresponding genome. Names highlighted in gray correspond to flies in the Shizophora group.

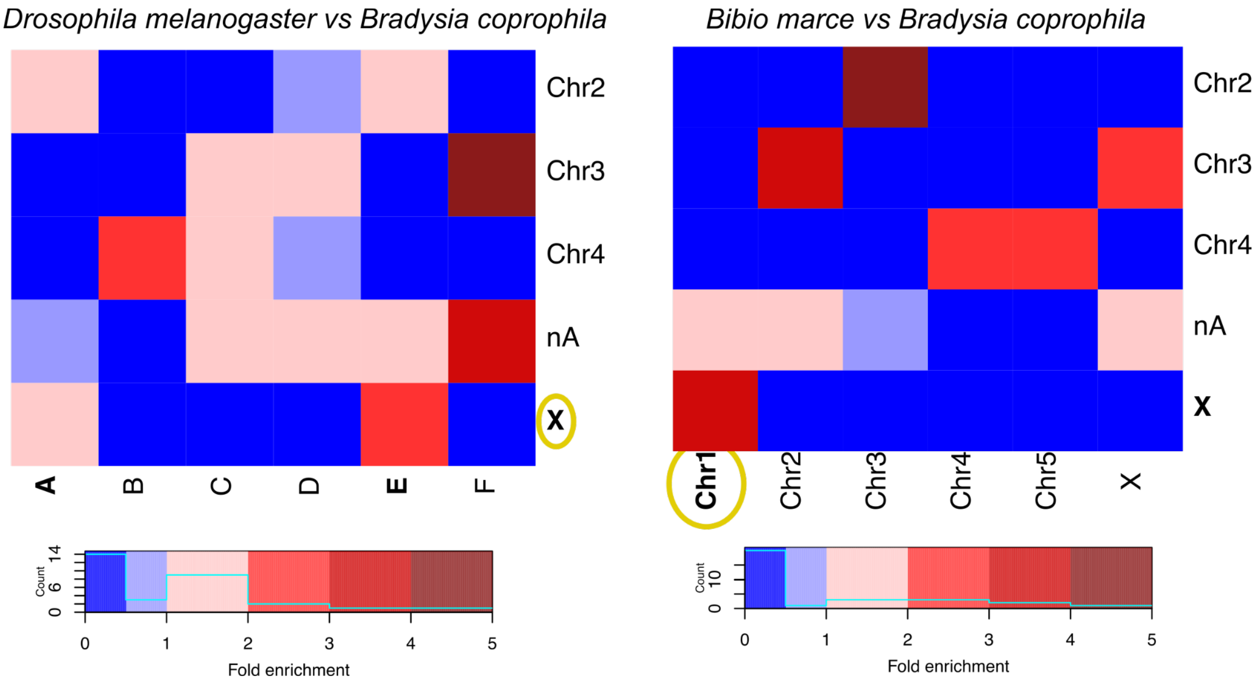

**Figure S2:** Heatmap showing the enrichment of Muller elements (x-axis) in the chromosomes of *Bradysia coprophila* (y-axis) (left) and the enrichment of *B. coprophila* chromosomes (x-axis) in the *Bibio marce* chromosomes (right). The X (elements A and E) in *B. coprophila* correspond to an autosome (Chr1) in the sister family Bibionidae (*Bibio marce)*, showing a simple turnover event.

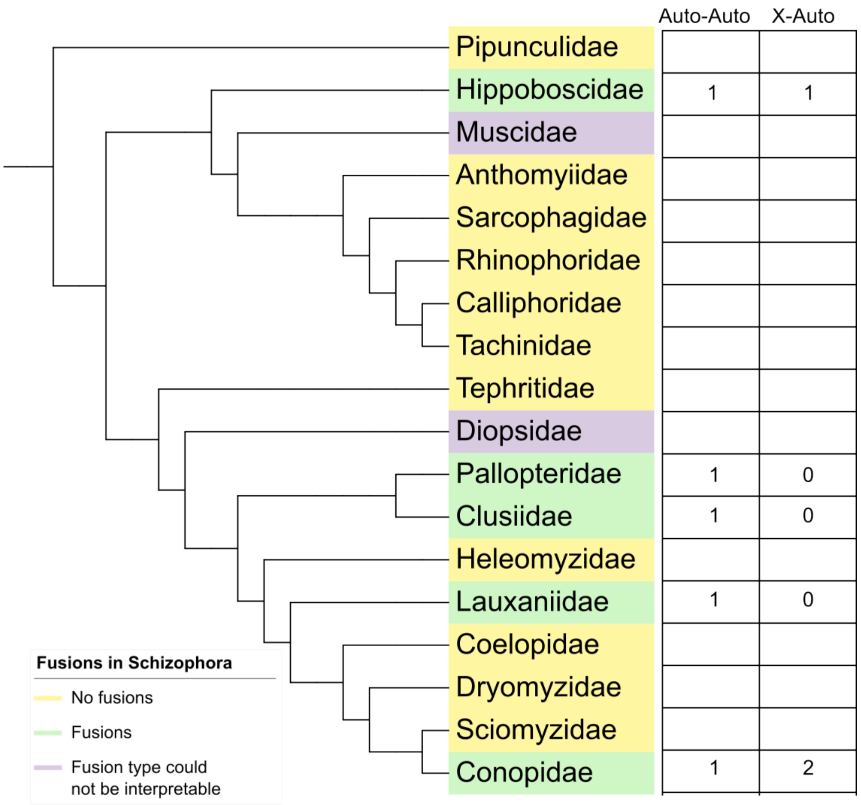

**Figure S3**: Fusions in the section Schizophora (where at least one chromosome-level assembly was available). Fusion type was not interpretable in Muscidae because X chromosomes are not characterized.

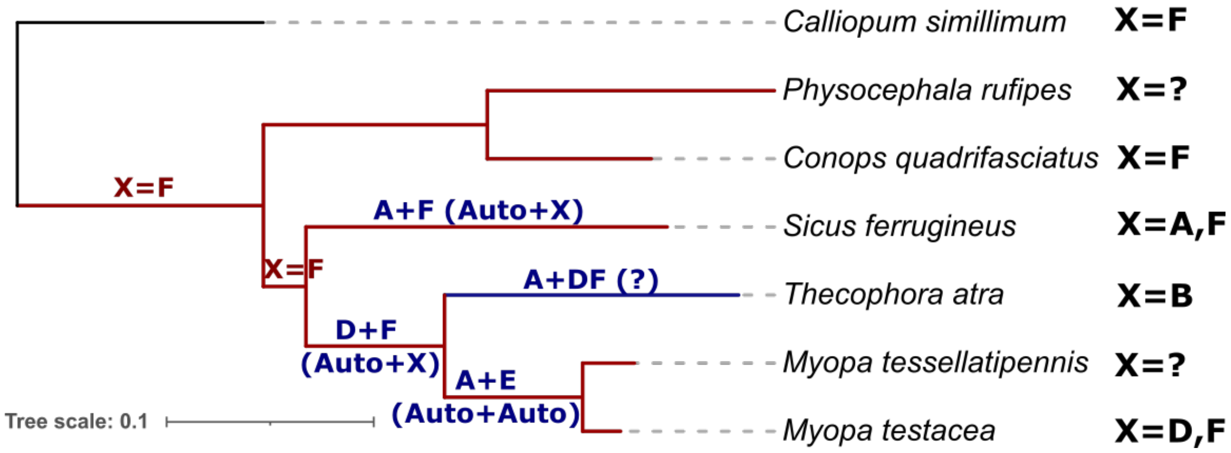

**Figure S4:** The X chromosome composition of Conopidae species and fusions involving the X and the autosomes. The red branches indicate that F has been maintained as the X, or even if another element was fused to it. The blue branch represents a turnover event.

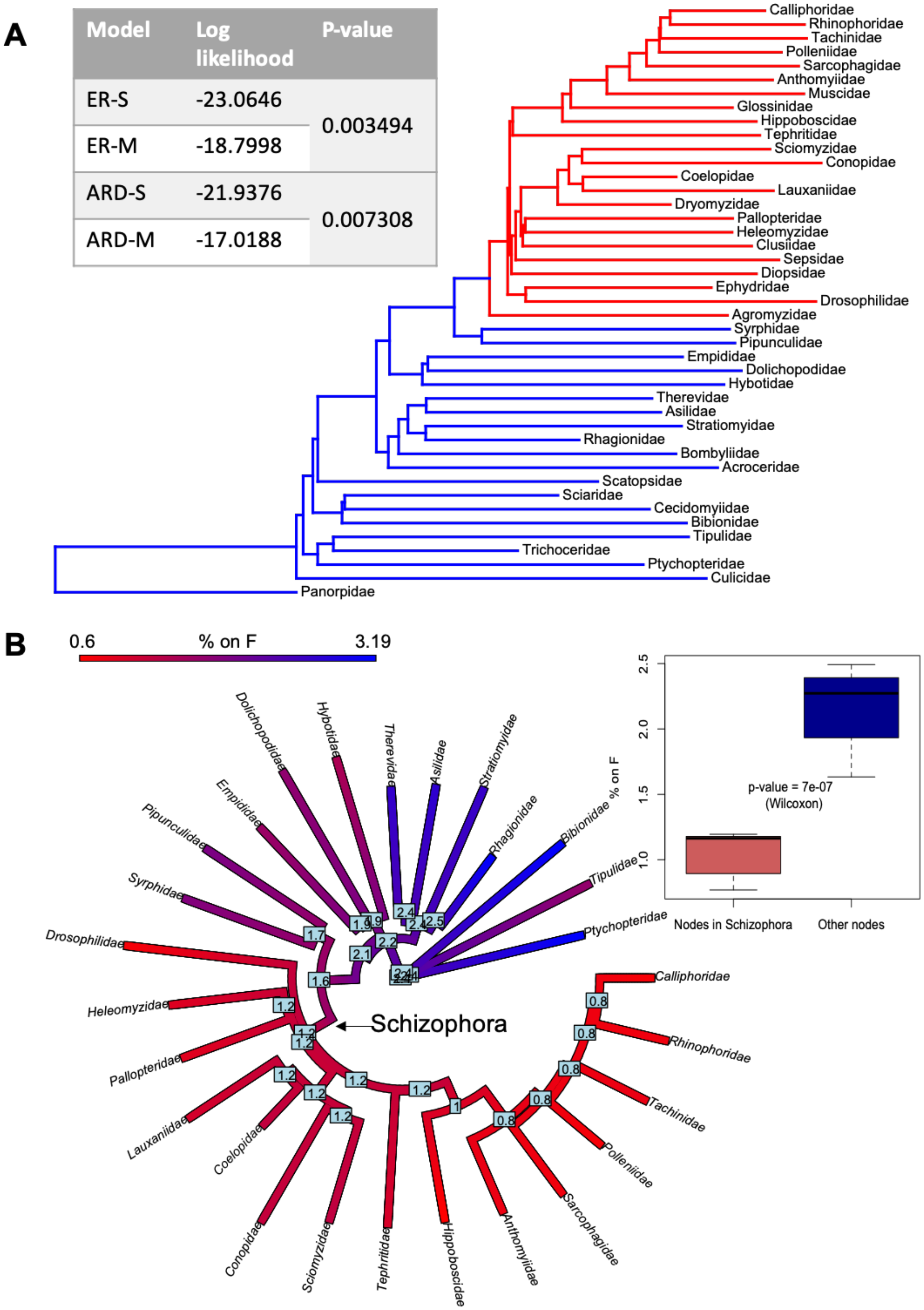

| Model | Log likelihood | P-value |
|---|---|---|
| ER-S | -23.0646 | 0.003494 |
| ER-M | -18.7998 | |
| ARD-S | -21.9376 | 0.007308 |
| ARD-M | -17.0188 | |

**Figure S5: Schizophora have increased turnover rates and a smaller proportion of genes on element F.** Full legend on the next page.

**Figure S5 (continued):** Panel A: Comparison of models with and without different turnover rates. The tree was split between Schizophora (red) and other branches (blue). The Phytools functions fitMk and fitmultiMk were used to compare the log-likelihood of models that assume a single turnover rate for the whole tree (ER-S and ARD-S) to models that assume two different turnover rates for the red and blue parts of the phylogeney (ER-M and ARD-M). ER assumes turnover and turnover reversal happen at equal rates, while ARD allows for asymmetric turnover and reveral rates. P-values were obtained with likelihood ratio tests. B. Ancestral reconstruction of the percentage of mapped genes located on element F throughout the dipteran phylogeny. Families for which only scaffold level assemblies were available, or where element F was fused to one of the large chromosomes, were excluded from this analysis (as it was not possible to infer the % of genes on F). Ancestral states were inferred with the Phytools function fastAnc. The labels at internal nodes show the inferred ancestral values for those nodes.

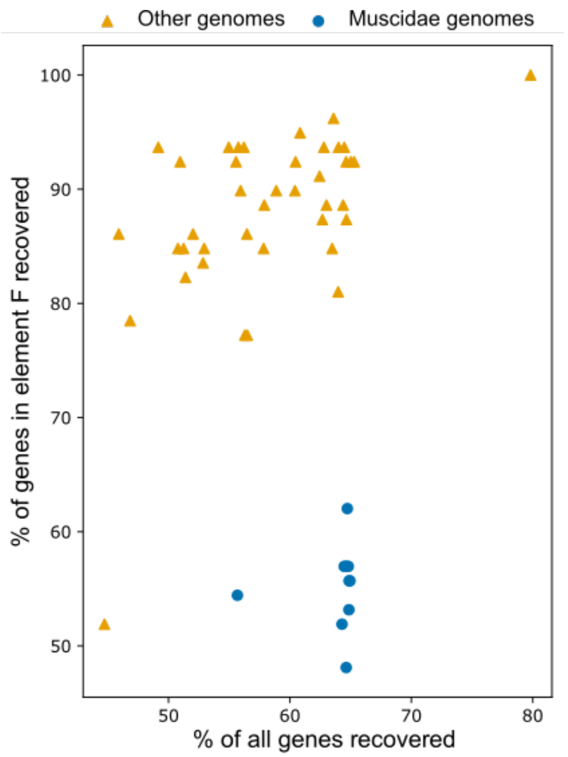

**Figure S6:** Fewer element F genes are recovered in Muscidae genomes (circles) than in other dipteran genomes (triangles) once divergence has been accounted for. The percentages refer to the percentage of *D. melanogaster* genes for which a mapping location was inferred in each of the diptera genomes, either when all genes are considered (x-axis) or when only element F genes are considered (y-axis).

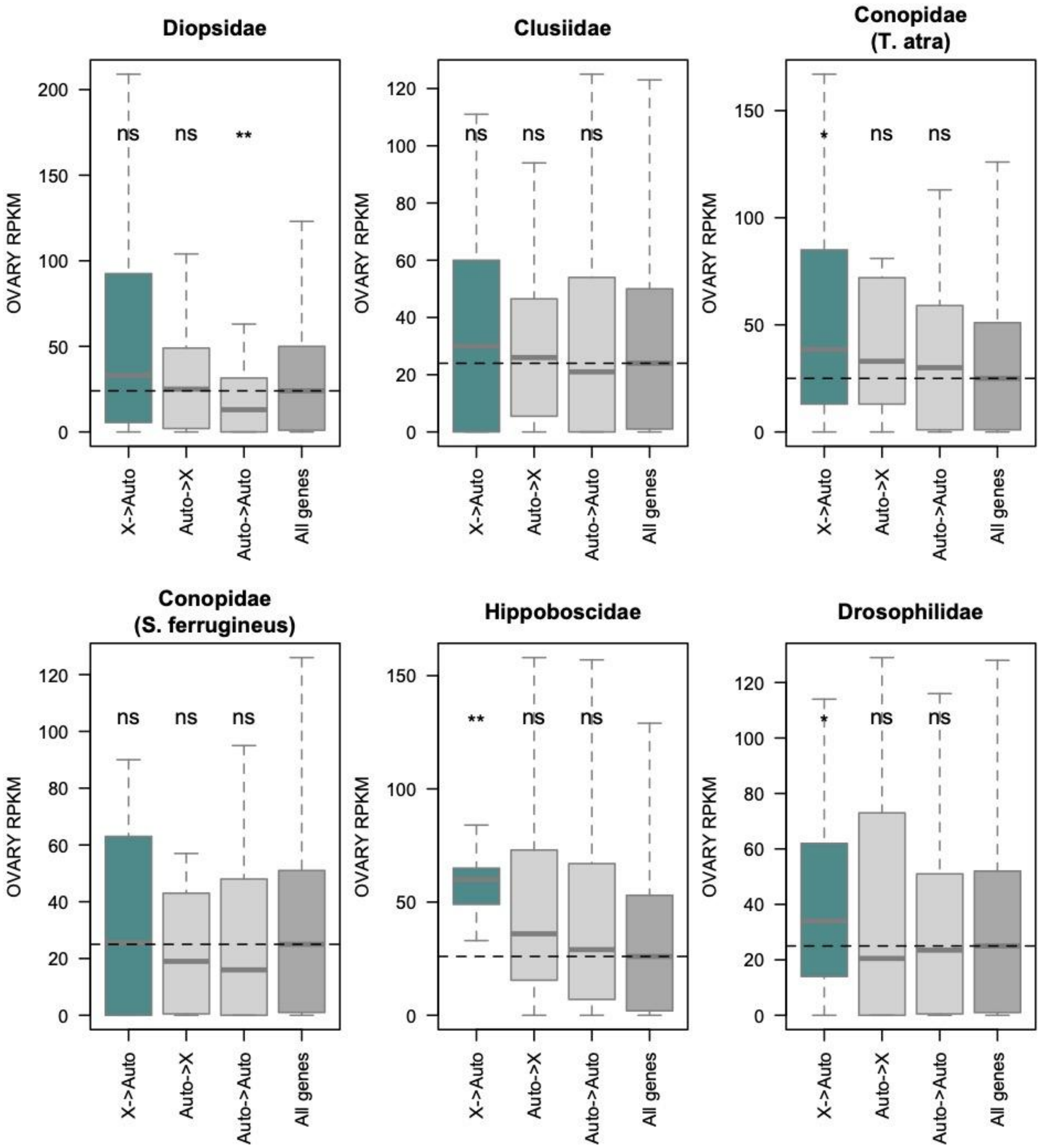

**Figure S7:** ***D. melanogaster*** **ovary expression of homologs inferred to have moved chromosomes, and that of all mapped genes, for each of the trios (named after the focal lineage under study).** X->Auto refers to genes that moved out of the X, Auto->X refers to genes that moved into the X, Auto->Auto refers to genes that moved between autosomes, and All genes refers to all mapped genes in each trio. Significance was assessed with Wilcoxon tests (* for p<0.05, ** for p<0.01, *** for p<0.001, ns for non significant).

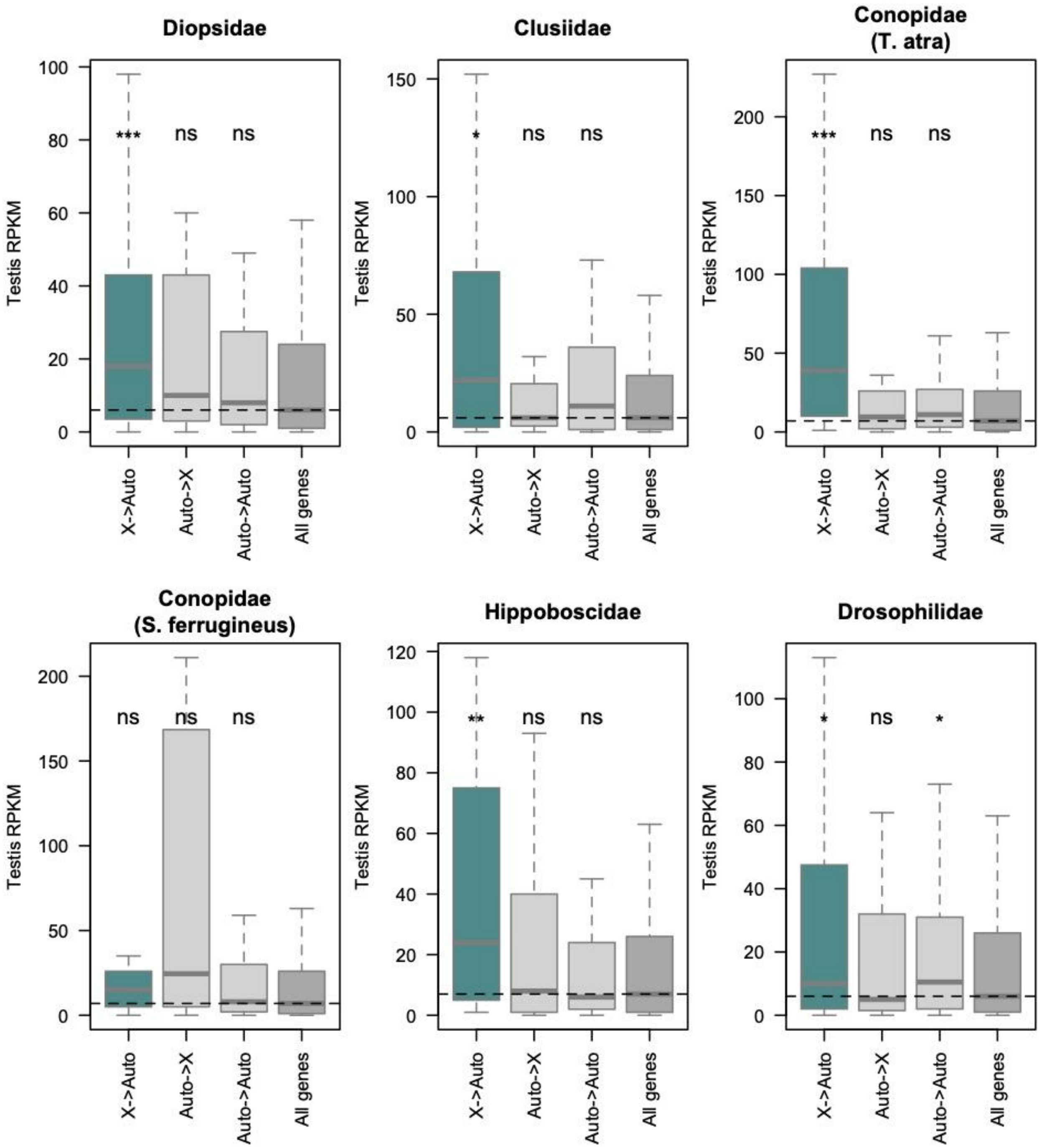

**Figure S8:** ***D. melanogaster*** **testis expression of homologs inferred to have moved chromosomes, and that of all mapped genes, for each of the trios (named after the focal lineage under study), using modENCODE data obtained from 4 day old mated males.** X->Auto refers to genes that moved out of the X, Auto->X refers to genes that moved into the X, Auto->Auto refers to genes that moved between autosomes, and All genes refers to all mapped genes in each trio. Significance was assessed with Wilcoxon tests (* for p<0.05, ** for p<0.01, *** for p<0.001, ns for non significant).

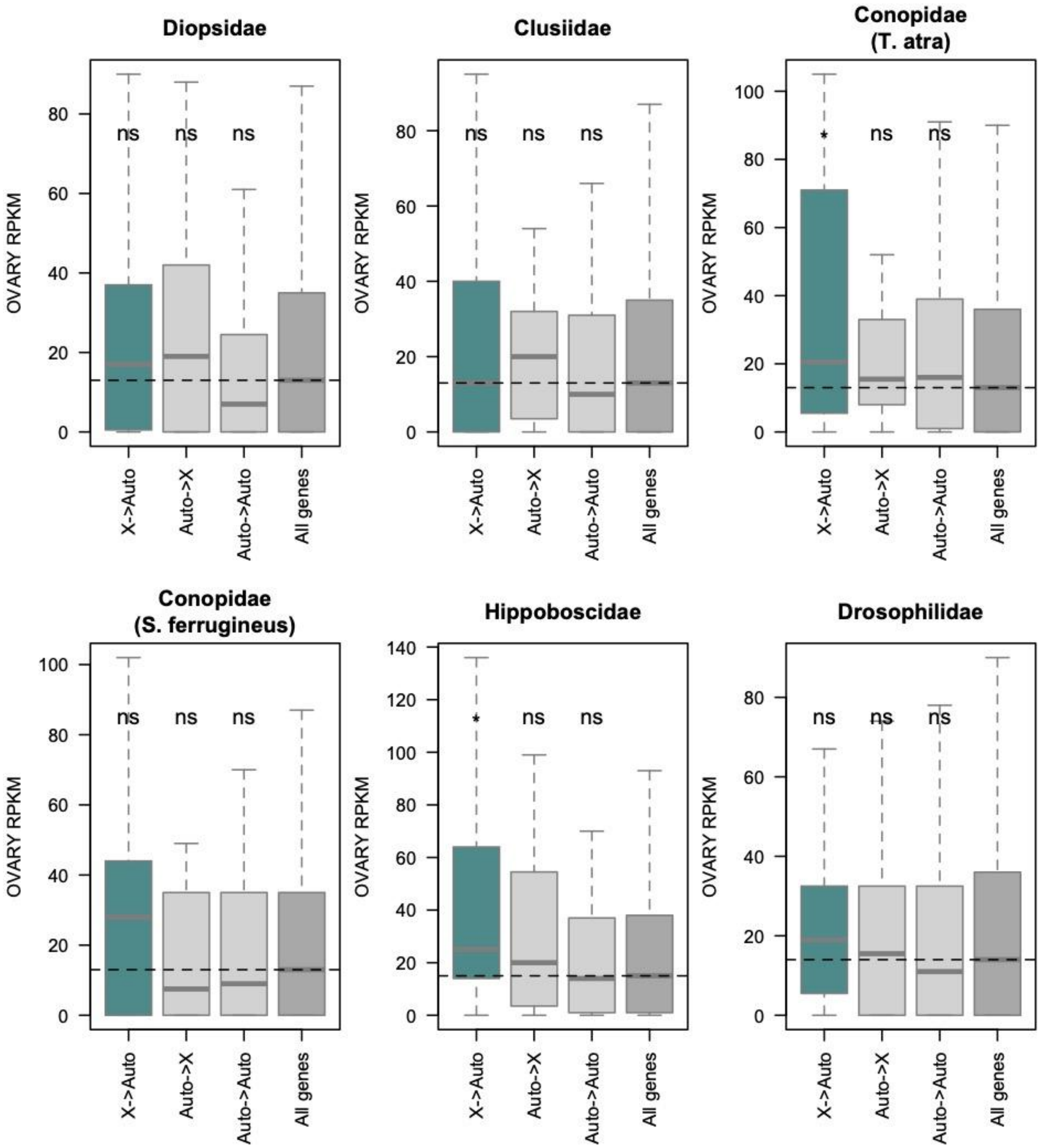

**Figure S9:** ***D. melanogaster*** **ovary expression of homologs inferred to have moved chromosomes, and that of all mapped genes, for each of the trios (named after the focal lineage under study), using modENCODE data obtained from 4 day old mated females.** X->Auto refers to genes that moved out of the X, Auto->X refers to genes that moved into the X, Auto->Auto refers to genes that moved between autosomes, and All genes refers to all mapped genes in each trio. Significance was assessed with Wilcoxon tests (* for p<0.05, ** for p<0.01, *** for p<0.001, ns for non significant).

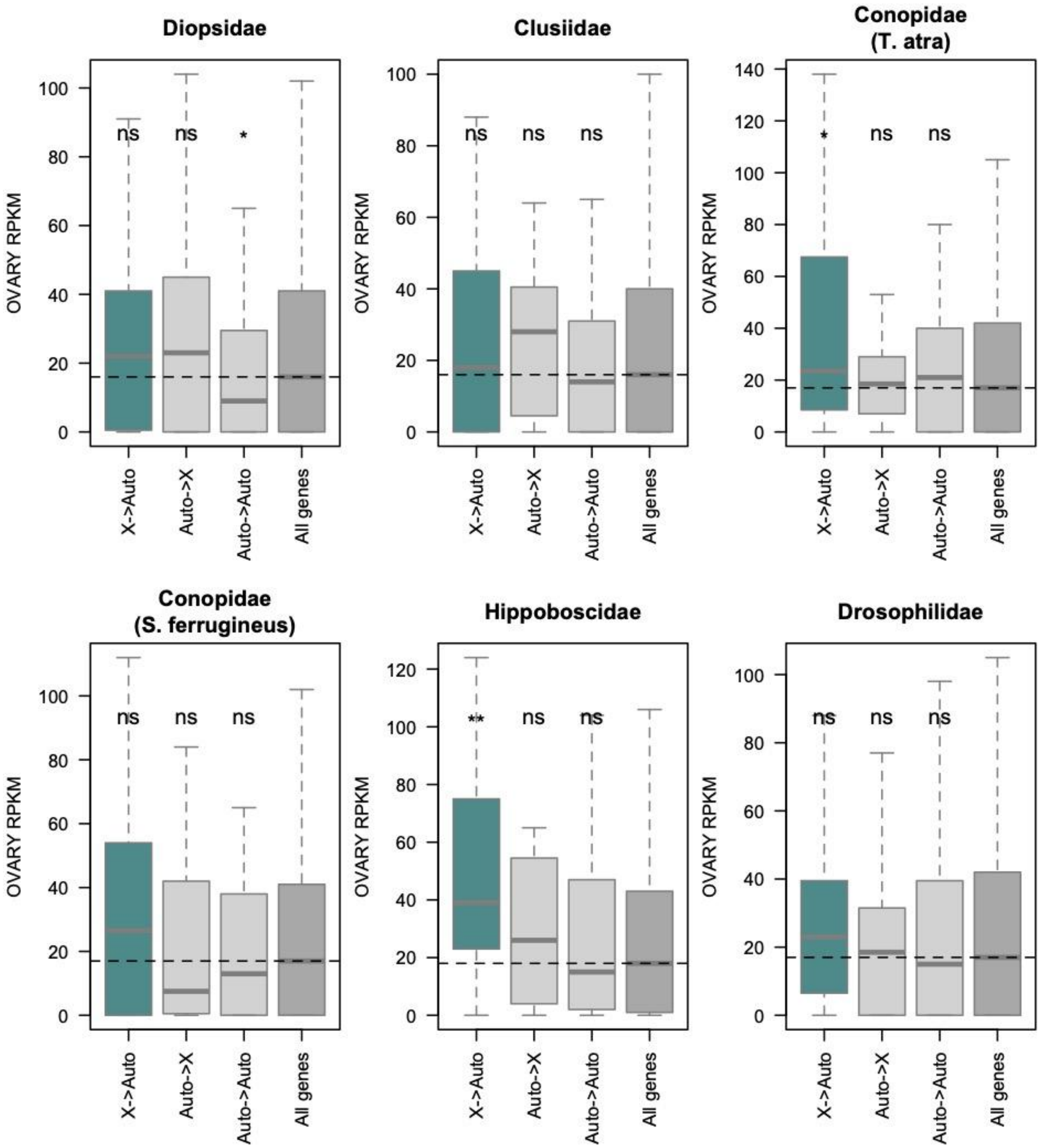

**Figure S10:** ***D. melanogaster*** **ovary expression of homologs inferred to have moved chromosomes, and that of all mapped genes, for each of the trios (named after the focal lineage under study), using modENCODE data obtained from 4 day old virgin females.** X->Auto refers to genes that moved out of the X, Auto->X refers to genes that moved into the X, Auto->Auto refers to genes that moved between autosomes, and All genes refers to all mapped genes in each trio. Significance was assessed with Wilcoxon tests (* for p<0.05, ** for p<0.01, *** for p<0.001, ns for non significant).

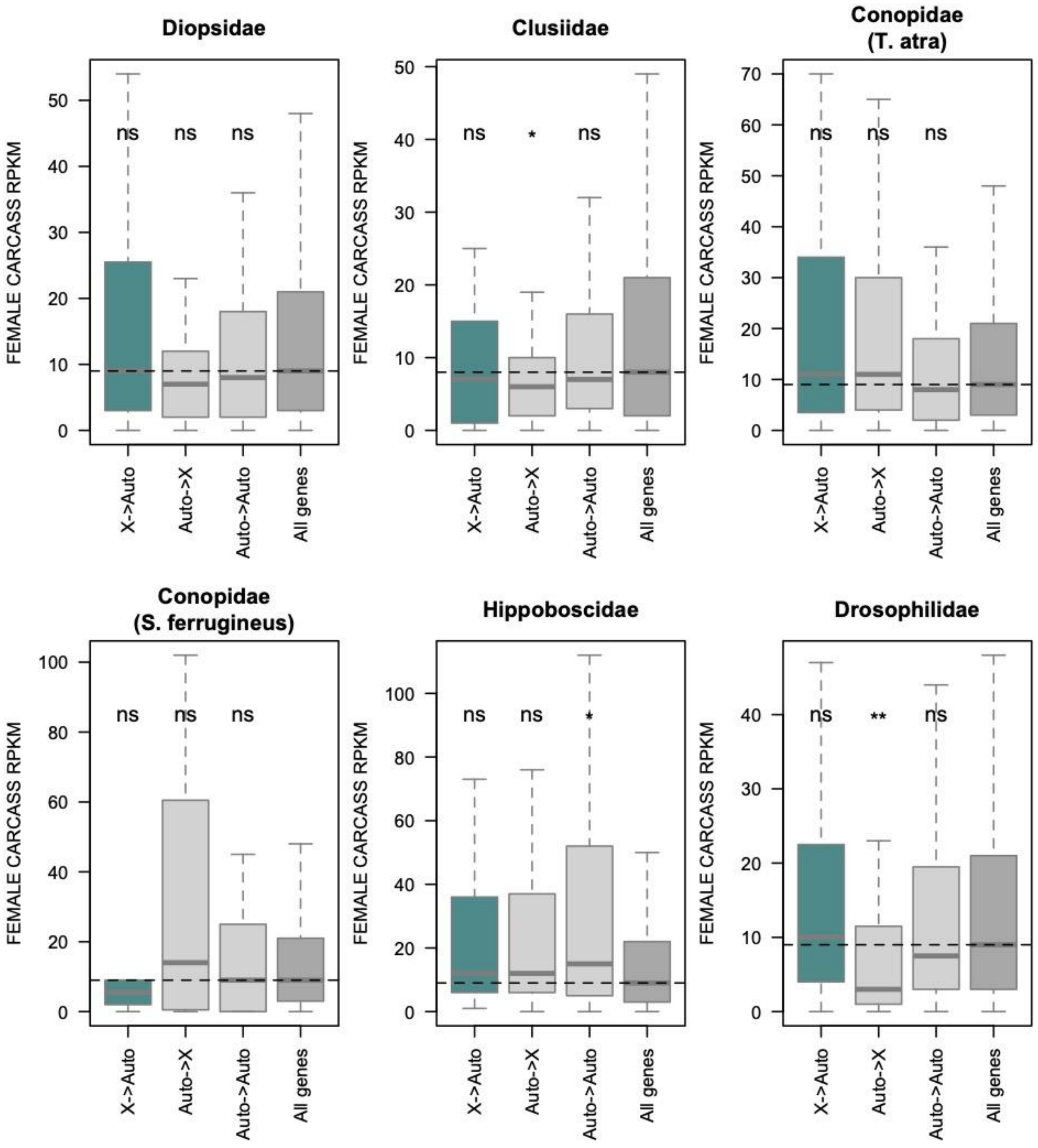

**Figure S11:** ***D. melanogaster*** **female carcass expression of homologs inferred to have moved chromosomes, and that of all mapped genes, for each of the trios (named after the focal lineage under study).** X->Auto refers to genes that moved out of the X, Auto->X refers to genes that moved into the X, Auto->Auto refers to genes that moved between autosomes, and All genes refers to all mapped genes in each trio. Significance was assessed with Wilcoxon tests (* for p<0.05, ** for p<0.01, *** for p<0.001, ns for non significant).

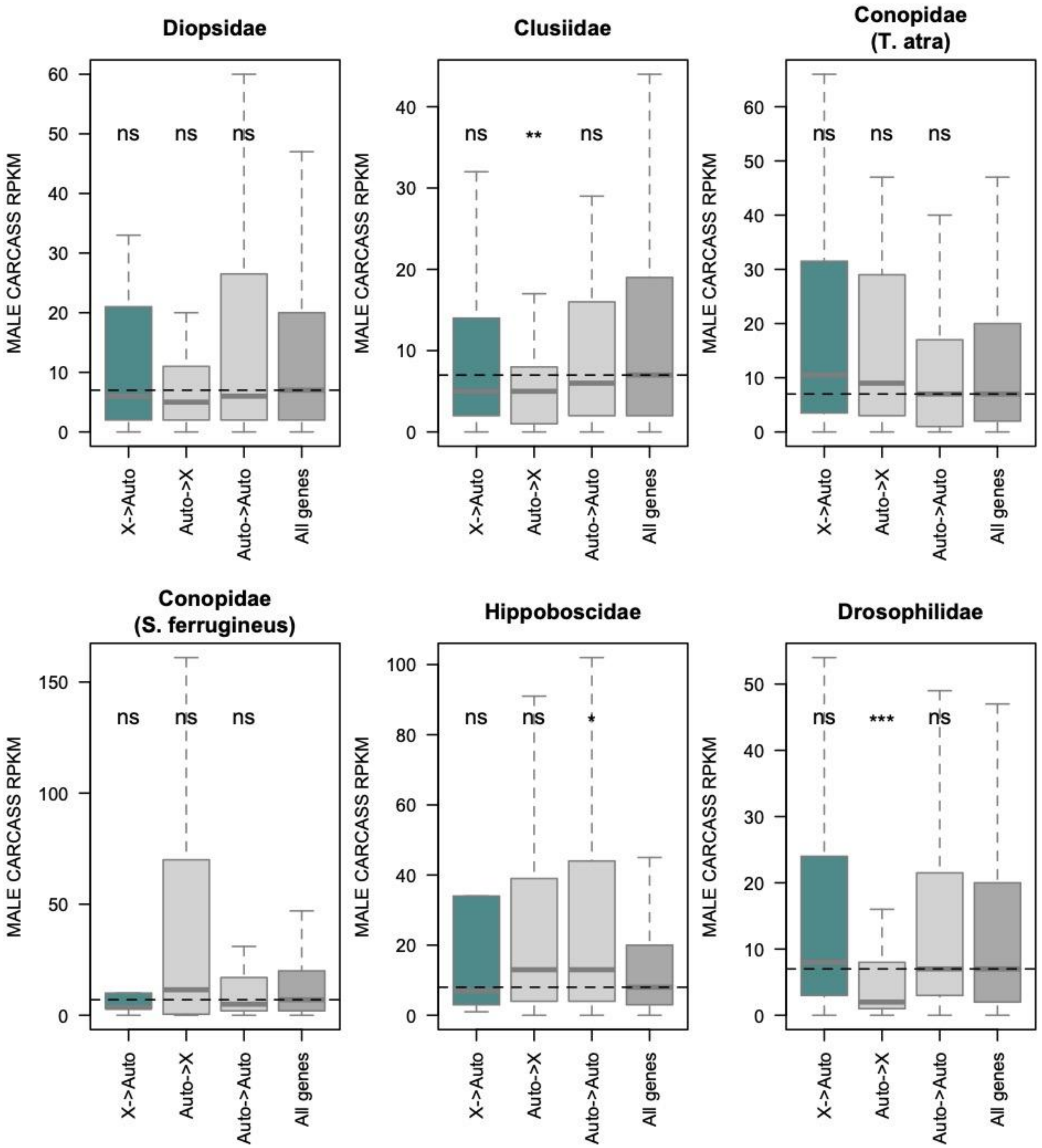

**Figure S12: *D. melanogaster* male carcass expression of homologs inferred to have moved chromosomes, and that of all mapped genes, for each of the trios (named after the focal lineage under study).** X->Auto refers to genes that moved out of the X, Auto->X refers to genes that moved into the X, Auto->Auto refers to genes that moved between autosomes, and All genes refers to all mapped genes in each trio. Significance was assessed with Wilcoxon tests (* for p<0.05, ** for p<0.01, *** for p<0.001, ns for non significant).

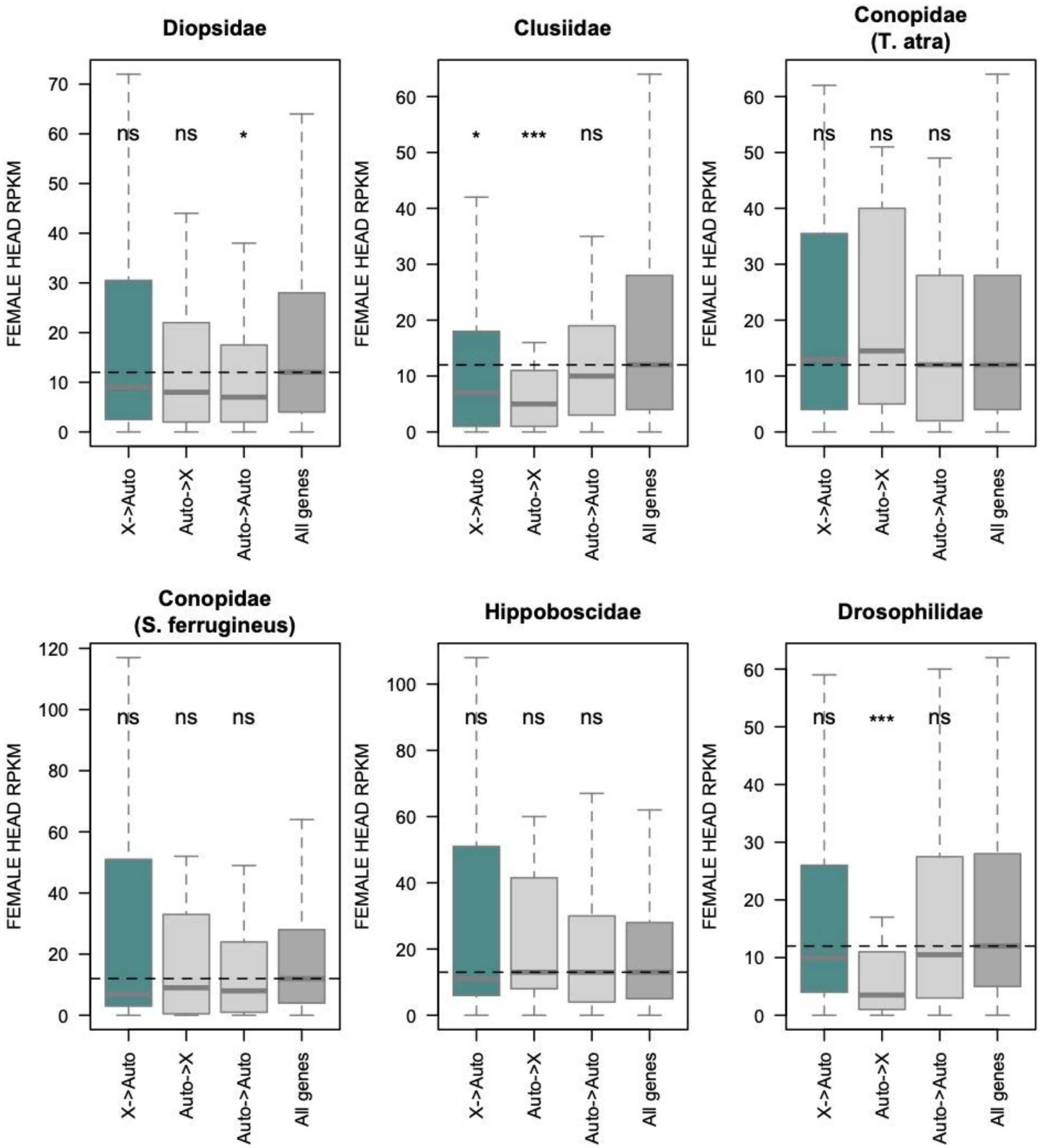

**Figure S13:** ***D. melanogaster*** **female head expression of homologs inferred to have moved chromosomes, and that of all mapped genes, for each of the trios (named after the focal lineage under study).** X->Auto refers to genes that moved out of the X, Auto->X refers to genes that moved into the X, Auto->Auto refers to genes that moved between autosomes, and All genes refers to all mapped genes in each trio. Significance was assessed with Wilcoxon tests (* for p<0.05, ** for p<0.01, *** for p<0.001, ns for non significant).

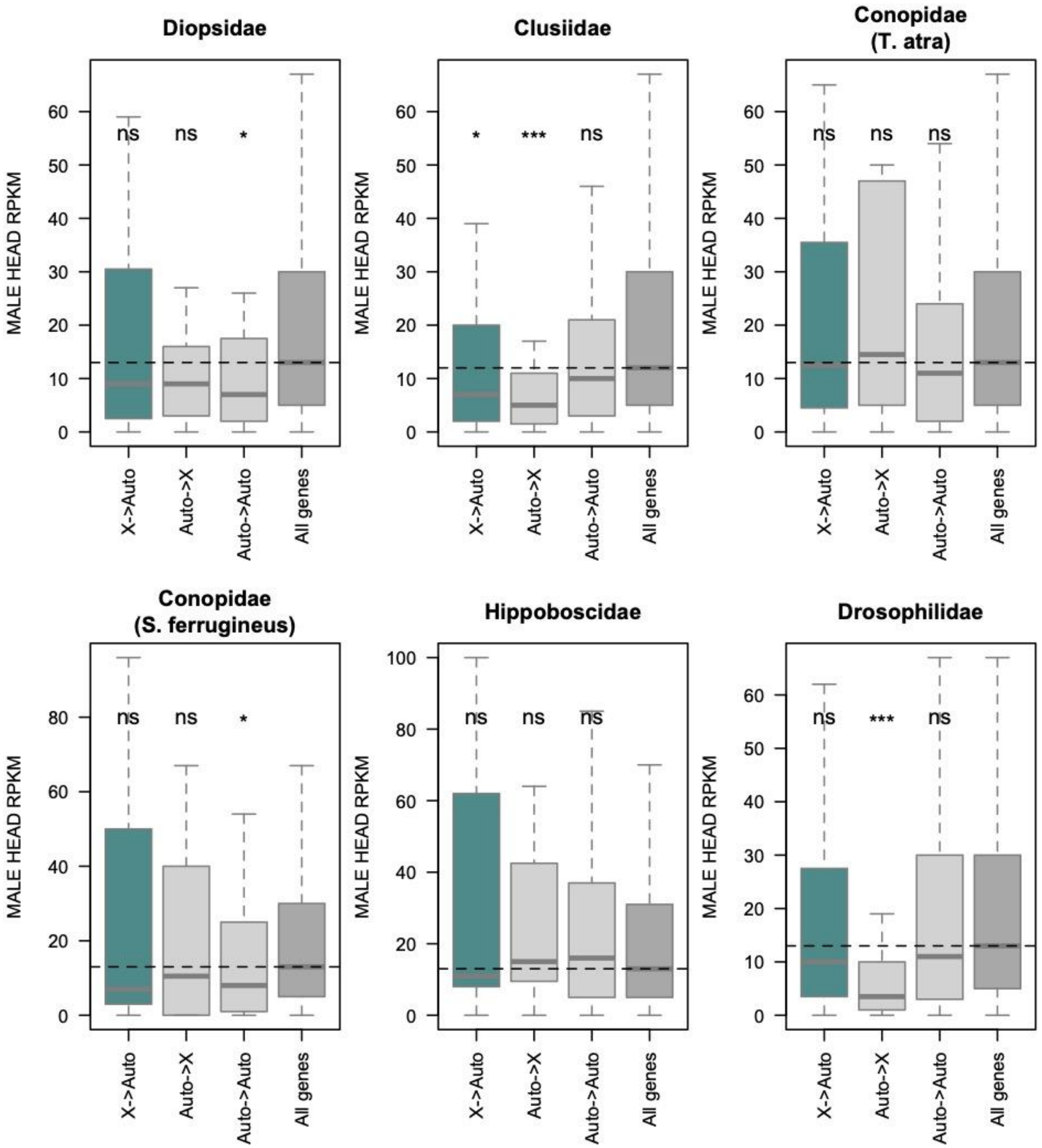

**Figure S14: *D. melanogaster* male head expression of homologs inferred to have moved chromosomes, and that of all mapped genes, for each of the trios (named after the focal lineage under study).** X->Auto refers to genes that moved out of the X, Auto->X refers to genes that moved into the X, Auto->Auto refers to genes that moved between autosomes, and All genes refers to all mapped genes in each trio. Significance was assessed with Wilcoxon tests (* for p<0.05, ** for p<0.01, *** for p<0.001, ns for non significant).

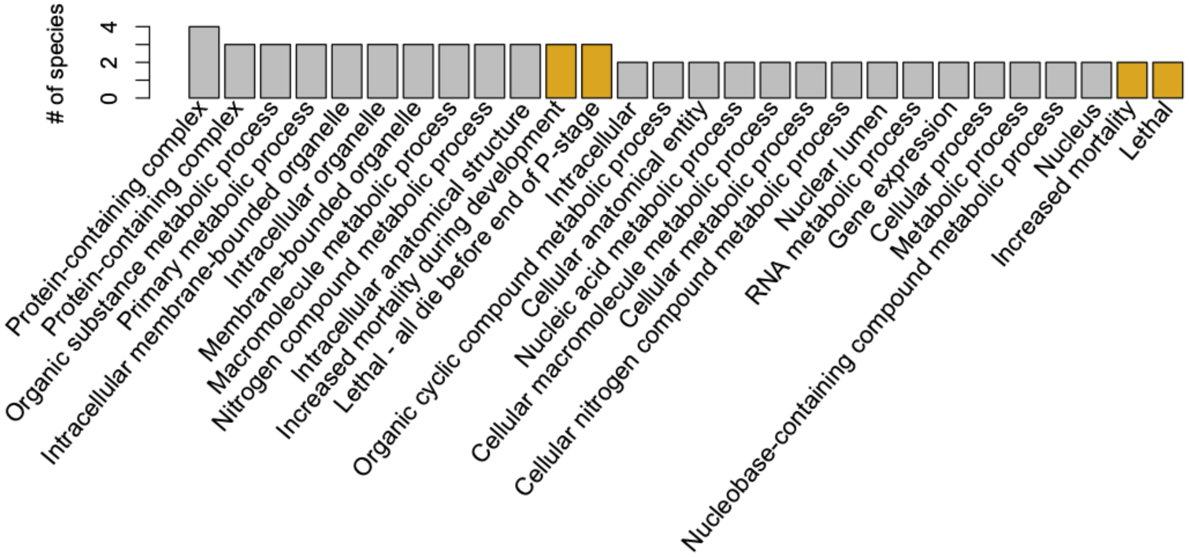

**Figure S15:** StringDB functional enrichment of genes that moved out of the X. The barplot shows the gene ontology (grey) and phenotype (yellow) terms that were enriched among genes that moved out of the X in at least 2 species, and the number of species this enrichment was found in.

**Supplementary Tables**

**Table S3:** The proportion of mapped genes found on the chromosome homologous to element F in species chosen to represent each of the families (shown in Figure 2). Lineages where element F is fused to another chromosome were excluded.

| **Species** | **Family** | **% of genes on F-homologue** |
|---|---|---|
| *Pegoplata infirma* | Anthomyiidae | 0.79% |
| *Dysmachus trigonus* | Asilidae | 2.73% |
| *Bibio marce* | Bibionidae | 3.18% |
| *Bellardia pandia* | Calliphoridae | 0.66% |
| *Clusia tigrina* | Clusiidae | 1.15% |
| *Coelopa pilipes* | Coelopidae | 1.23% |
| *Conops quadrifasciatus* | Conopidae | 1.21% |
| *Dolichopus griseipennis* | Dolichopodidae | 1.73% |
| *Drosophila pseudoobscura* | Drosophilidae | 0.70% |
| *Dryomyza anilis* | Dryomyzidae | 1.02% |
| *Rhamphomyia laevipes* | Empididae | 1.62% |
| *Suillia variegata* | Heleomyzidae | 0.93% |
| *Ornithomya chloropus* | Hippoboscidae | 0.60% |
| *Hybos culiciformis* | Hybotidae | 1.45% |
| *Calliopum simillimum* | Lauxaniidae | 1.03% |
| *Toxonevra muliebris* | Pallopteridae | 0.90% |
| *Nephrocerus scutellatus* | Pipunculidae | 1.94% |
| *Pollenia amentaria* | Polleniidae | 0.68% |
| *Ptychoptera albimana* | Ptychopteridae | 3.19% |
| *Rhagio lineola* | Rhagionidae | 3.15% |
| *Melanophora roralis* | Rhinophoridae | 0.74% |
| *Sarcophaga caerulescens* | Sarcophagidae | 0.74% |
| *Coremacera marginata* | Sciomyzidae | 1.25% |
| *Beris chalybata* | Stratiomyidae | 2.64% |
| *Blera fallax* | Syrphidae | 1.87% |
| *Cistogaster globosa* | Tachinidae | 0.80% |
| *Anastrepha ludens* | Tephritidae | 1.01% |

| | | |
|---|---|---|
| *Thereva nobilitata* | Therevidae | 2.67% |
| *Nephrotoma appendiculata* | Tipulidae | 1.69% |

**Table S4: Protein annotations used for our Diptera-branched tree.**

| Busco | Family | *Species* | Link to annotations |
|---|---|---|---|
| | Acroceridae | *Acrocera orbiculus* | https://ftp.ebi.ac.uk/pub/ensemblorganisms/Acrocera_orbiculus/GCA_947359355.1/ensembl/geneset/2023_05/ |
| | Anthomyiidae | *Pegoplata infirma* | https://ftp.ebi.ac.uk/pub/ensemblorganisms/Pegoplata_infirma/GCA_963921195.1/braker/geneset/2024_05/ |
| | Asilidae | *Machimus atricapillus* | https://ftp.ebi.ac.uk/pub/ensemblorganisms/Machimus_atricapillus/GCA_933228815.1/ensembl/geneset/2022_08/ |
| | Bibionidae | *Bibio marci* | https://ftp.ebi.ac.uk/pub/ensemblorganisms/Bibio_marci/GCA_910594885.2/ensembl/geneset/2024_05/ |
| | Bombyliidae | *Bombylius discolor* | https://ftp.ebi.ac.uk/pub/ensemblorganisms/Bombylius_discolor/GCA_939192795.1/ensembl/geneset/2022_07/ |
| | Calliphoridae | *Bellardia pandia* | https://ftp.ebi.ac.uk/pub/ensemblorganisms/Bellardia_pandia/GCA_916048285.2/braker/geneset/2022_03/ |
| | Clusiidae | *Clusia tigrina* | https://ftp.ebi.ac.uk/pub/ensemblorganisms/Clusia_tigrina/GCA_920105625.2/braker/geneset/2023_07/ |
| BUSCO | Coelopidae | *Coelopa pilipes* | GCA_947389925.1 |
| | Conopidae | *Thecophora atra* | https://ftp.ebi.ac.uk/pub/ensemblorganisms/Thecophora_atra/GCA_937620795.1/braker/geneset/2022_07/ |
| | Culicidae | *Anopheles gambiae* | GCF_943734735.2 |
| | Diopsidae | *Teleopsis dalmanni* | GCF_002237135.1 |
| | Dolichopodidae | *Poecilobothrus nobilitatus* | https://ftp.ebi.ac.uk/pub/ensemblorganisms/Poecilobothrus_nobilitatus/GCA_947095535.1/ensembl/geneset/2024_05/ |
| | Drosophilidae | *Drosophila melanogaster* | GCF_000001215.4 |
| BUSCO | Dryomyzidae | *Dryomyza anilis* | GCA_951804985.1 |
| | Empididae | *Empis livida* | https://ftp.ebi.ac.uk/pub/ensemblorganisms/Empis_livida/GCA_963932195.1/ensembl/geneset/2024_05/ |
| | Heleomyzidae | *Suillia variegata* | https://ftp.ebi.ac.uk/pub/ensemblorganisms/Suillia_variegata/GCA_949127995.1/braker/geneset/2023_07/ |
| BUSCO | Hippoboscidae | *Ornithomya fringillina* | GCA_963978525.1 |
| | Hybotidae | *Hybos culiciformis* | https://ftp.ebi.ac.uk/pub/ensemblorganisms/Hybos_culic |

| | | | |
|---|---|---|---|
| | | | iformis/GCA_964007475.1/braker/geneset/2024_05/ |
| | Lauxaniidae | *Calliopum simillimum* | https://ftp.ebi.ac.uk/pub/ensemblorganisms/Calliopum_simillimum/GCA_951812925.1/braker/geneset/2023_10/ |
| | Muscidae | *Hydrotaea cyrtoneurina* | https://ftp.ebi.ac.uk/pub/ensemblorganisms/Hydrotaea_cyrtoneurina/GCA_958296145.1/ensembl/geneset/2024_05/ |
| | Pallopteridae | *Toxonevra muliebris* | https://ftp.ebi.ac.uk/pub/ensemblorganisms/Toxonevra_muliebris/GCA_963691655.1/braker/geneset/2024_05/ |
| | Pipunculidae | *Nephrocerus scutellatus* | https://ftp.ebi.ac.uk/pub/ensemblorganisms/Nephrocerus_scutellatus/GCA_947095585.1/ensembl/geneset/2024_04/ |
| | Polleniidae | *Pollenia amentaria* | https://ftp.ebi.ac.uk/pub/ensemblorganisms/Pollenia_amentaria/GCA_943735925.1/ensembl/geneset/2022_08/ |
| | Ptychopteridae | *Ptychoptera contaminata* | https://ftp.ebi.ac.uk/pub/ensemblorganisms/Ptychoptera_contaminata/GCA_963942525.1/ensembl/geneset/2024_05/ |
| BUSCO | Rhagionidae | *Rhagio lineola* | GCA_964211845.1 |
| | Rhinophoridae | *Phyto melanocephala* | https://ftp.ebi.ac.uk/pub/ensemblorganisms/Phyto_melanocephala/GCA_941918925.1/braker/geneset/2022_07/ |
| | Sarcophagidae | *Sarcophaga caerulescens* | https://ftp.ebi.ac.uk/pub/ensemblorganisms/Sarcophaga_caerulescens/GCA_927399465.1/ensembl/geneset/2022_03/ |
| | Sciaridae | *Pseudolycoriella hygida* | GCA_029228625.1 |
| | Sciomyzidae | *Coremacera marginata* | https://ftp.ebi.ac.uk/pub/ensemblorganisms/Coremacera_marginata/GCA_914767935.1/ensembl/geneset/2021_12/ |
| | Stratiomyidae | *Chorisops tibialis* | https://ftp.ebi.ac.uk/pub/ensemblorganisms/Chorisops_tibialis/GCA_963669355.1/braker/geneset/2024_05/ |
| | Syrphidae | *Blera fallax* | https://ftp.ebi.ac.uk/pub/ensemblorganisms/Blera_fallax/GCA_946965025.1/ensembl/geneset/2024_04/ |
| | Tachinidae | *Cistogaster globosa* | https://ftp.ebi.ac.uk/pub/ensemblorganisms/Cistogaster_globosa/GCA_937654795.1/braker/geneset/2022_07/ |
| | Tephritidae | *Anastrepha ludens* | GCF_028408465.1 |
| | Therevidae | *Thereva nobiliata* | https://ftp.ebi.ac.uk/pub/ensemblorganisms/Thereva_nobilitata/GCA_963855945.1/braker/geneset/2024_05/ |
| | Tipulidae | *Nephrotoma appendiculata* | https://ftp.ebi.ac.uk/pub/ensemblorganisms/Nephrotoma_appendiculata/GCA_947310385.1/ensembl/geneset/ |

| | | | |
|---|---|---|---|
| | | | 2023_05/ |
| | Sepsidae | *Orygma luctuosum* | https://ftp.ebi.ac.uk/pub/ensemblorganisms/Orygma_luctuosum/GCA_965218485.1/ensembl/geneset/2025_04/ |
| | Panorpidae | *Panorpa germanica* | https://ftp.ebi.ac.uk/pub/ensemblorganisms/Panorpa_germanica/GCA_963678705.1/ |
| **Scaffolded assemblies:** | | | |
| BUSCO | Glossinidae | *Glossina fuscipes* | GCF_014805625.2 |
| BUSCO | Ephydridae | *Cirrula hians* | |
| BUSCO | Agromyzidae | *Liriomyza trifolii* | GCA_043165165.1 |
| BUSCO | Cecidomyiidae | *Sitodiplosis mosellanafa* | GCF_009176505.1 |
| BUSCO | Scatopsidae | *Coboldia fuscipes* | GCA_001014335.1 |
| BUSCO | Trichoceridae | *Trichoceridae species* | GCA_001014425.1 |

**Table S7:** GO enrichment amount genes that moved out of the X in Diopsidae. Rows highlighted in yellow show GO terms that remain enriched when mapped genes are used as the background to test against.

| #category | term ID | term description | observed gene count | false discovery rate |
|---|---|---|---|---|
| GO Component | GO:0032991 | Protein-containing complex | 30 | 0.0019 |
| GO Component | GO:0005622 | Intracellular anatomical structure | 48 | 0.003 |
| GO Component | GO:0043226 | Organelle | 43 | 0.005 |
| GO Component | GO:0043227 | Membrane-bounded organelle | 40 | 0.0058 |
| GO Component | GO:0140535 | Intracellular protein-containing complex | 11 | 0.0066 |
| GO Component | GO:0005665 | RNA polymerase II, core complex | 3 | 0.0095 |
| GO Component | GO:0005634 | Nucleus | 27 | 0.0106 |
| GO Component | GO:0043229 | Intracellular organelle | 41 | 0.0106 |
| GO Component | GO:0070013 | Intracellular organelle lumen | 14 | 0.0153 |
| GO Component | GO:0043231 | Intracellular membrane-bounded organelle | 37 | 0.0168 |
| GO Component | GO:0031981 | Nuclear lumen | 12 | 0.0195 |
| GO Component | GO:0005929 | Cilium | 6 | 0.0202 |
| GO Component | GO:1990234 | Transferase complex | 9 | 0.0305 |
| GO Component | GO:0070776 | MOZ/MORF histone acetyltransferase complex | 2 | 0.0317 |
| **KEGG** | **dme03020** | **RNA polymerase** | **4** | **0.0019** |

| | | | | |
|---|---|---|---|---|
| Reactome | DME-6781823 | Formation of TC-NER Pre-Incision Complex | 5 | 0.0048 |
| Reactome | DME-6782210 | Gap-filling DNA repair synthesis and ligation in TC-NER | 4 | 0.0342 |
| Reactome | DME-6782135 | Dual incision in TC-NER | 4 | 0.0343 |
| Reactome | DME-3700989 | Transcriptional Regulation by TP53 | 6 | 0.0465 |
| WikiPathways | WP177 | Eukaryotic transcription initiation | 3 | 0.0131 |
| WikiPathways | WP406 | Mitochondrial long chain fatty acid beta-oxidation | 2 | 0.015 |
| COMPARTMENTS | GOCC:0005665 | RNA polymerase II, core complex | 3 | 0.0203 |
| COMPARTMENTS | GOCC:0005929 | Cilium | 6 | 0.0203 |
| COMPARTMENTS | GOCC:0031981 | Nuclear lumen | 12 | 0.0203 |
| COMPARTMENTS | GOCC:0032991 | Protein-containing complex | 28 | 0.0203 |
| COMPARTMENTS | GOCC:0043226 | Organelle | 34 | 0.0203 |
| COMPARTMENTS | GOCC:0070013 | Intracellular organelle lumen | 14 | 0.0203 |
| COMPARTMENTS | GOCC:0043229 | Intracellular organelle | 32 | 0.0283 |
| COMPARTMENTS | GOCC:0005622 | Intracellular | 37 | 0.0313 |
| UniProt Keywords | KW-0539 | Nucleus | 17 | 0.0068 |
| UniProt Keywords | KW-0240 | DNA-directed RNA polymerase | 3 | 0.0218 |
| InterPro | IPR036603 | RNA polymerase, RBP11-like subunit | 3 | 0.0122 |

**Table S8:** GO enrichment amount genes that moved out of the X in Clusiidae.

| #category | term ID | term description | observed gene count | false discovery rate |
|---|---|---|---|---|
| GO Process | GO:0006807 | Nitrogen compound metabolic process | 29 | 0.0072 |
| GO Process | GO:0010467 | Gene expression | 16 | 0.0072 |
| GO Process | GO:0043170 | Macromolecule metabolic process | 27 | 0.0072 |
| GO Process | GO:0044238 | Primary metabolic process | 30 | 0.0072 |
| GO Process | GO:0071704 | Organic substance metabolic process | 32 | 0.0072 |
| GO Process | GO:1901360 | Organic cyclic compound metabolic process | 17 | 0.0199 |
| GO Process | GO:0006725 | Cellular aromatic compound metabolic process | 16 | 0.043 |
| GO Component | GO:0032991 | Protein-containing complex | 23 | 0.0408 |
| Monarch | FBcv:0002000 | Lethal - all die before end of P-stage | 24 | 0.0061 |
| Monarch | FBcv:0002019 | Increased mortality during development | 31 | 0.0288 |
| COMPARTMENTS | GOCC:0032991 | Protein-containing complex | 26 | 0.0017 |

**Table S9:** GO enrichment amount genes that moved out of the X in Conopidae (*Thecophora atra*).

| #category | term ID | term description | observed gene count | false discovery rate |
|---|---|---|---|---|
| GO Process | GO:0044237 | Cellular metabolic process | 29 | 0.00054 |
| GO Process | GO:0008152 | Metabolic process | 31 | 0.0027 |
| GO Process | GO:0043170 | Macromolecule metabolic process | 26 | 0.0027 |
| GO Process | GO:0044260 | Cellular macromolecule metabolic process | 17 | 0.0027 |
| GO Process | GO:0090304 | Nucleic acid metabolic process | 15 | 0.0027 |
| GO Process | GO:0006807 | Nitrogen compound metabolic process | 27 | 0.0036 |
| GO Process | GO:0009987 | Cellular process | 39 | 0.0036 |
| GO Process | GO:0006139 | Nucleobase-containing compound metabolic process | 16 | 0.0043 |
| GO Process | GO:0034641 | Cellular nitrogen compound metabolic process | 18 | 0.0064 |
| GO Process | GO:0071704 | Organic substance metabolic process | 28 | 0.0112 |
| GO Process | GO:0044085 | Cellular component biogenesis | 15 | 0.0114 |
| GO Process | GO:0044238 | Primary metabolic process | 27 | 0.0114 |
| GO Process | GO:0071840 | Cellular component organization or biogenesis | 22 | 0.014 |
| GO Process | GO:0016571 | Histone methylation | 4 | 0.0155 |
| GO Process | GO:0022613 | Ribonucleoprotein complex biogenesis | 7 | 0.0183 |
| GO Process | GO:0016070 | RNA metabolic process | 11 | 0.0276 |

| | | | | |
|---|---|---|---|---|
| GO Process | GO:0042254 | Ribosome biogenesis | 6 | 0.0276 |
| GO Component | GO:0005622 | Intracellular anatomical structure | 39 | 8.84E-05 |
| GO Component | GO:0043229 | Intracellular organelle | 34 | 0.0011 |
| GO Component | GO:0043227 | Membrane-bounded organelle | 31 | 0.0045 |
| GO Component | GO:0043231 | Intracellular membrane-bounded organelle | 30 | 0.0058 |
| GO Component | GO:0005634 | Nucleus | 21 | 0.0164 |
| GO Component | GO:0032991 | Protein-containing complex | 21 | 0.0164 |
| GO Component | GO:0031981 | Nuclear lumen | 10 | 0.0255 |
| GO Component | GO:0110165 | Cellular anatomical entity | 41 | 0.0255 |
| GO Component | GO:0035097 | Histone methyltransferase complex | 3 | 0.0462 |
| Reactome | DME-6791226 | Major pathway of rRNA processing in the nucleolus and cytosol | 5 | 0.013 |
| Monarch | FBcv:0000351 | Lethal | 24 | 0.0383 |
| Monarch | FBcv:0000379 | Enhancer of variegation | 3 | 0.0383 |
| Monarch | FBcv:0000425 | Increased cell death | 8 | 0.0383 |
| Monarch | FBcv:0002000 | Lethal - all die before end of P-stage | 20 | 0.0383 |
| Monarch | FBcv:0002004 | Increased mortality | 30 | 0.0383 |
| Monarch | FBcv:0002019 | Increased mortality during development | 29 | 0.0383 |

| | | | | |
|---|---|---|---|---|
| Monarch | FBcv:0002027 | Lethal - all die before end of pupal stage | 19 | 0.0383 |
| UniProt Keywords | KW-0853 | WD repeat | 6 | 0.0074 |
| SMART | SM00320 | WD40 repeats | 6 | 0.0207 |

**Table S10:** GO enrichment amount genes that moved out of the X in Drosophilidae. Rows highlighted in yellow show GO terms that remain enriched when mapped genes are used as the background to test against.

| #category | term ID | term description | observed gene count | false discovery rate |
|---|---|---|---|---|
| **GO Process** | **GO:0006139** | **Nucleobase-containing compound metabolic process** | **33** | **1.69E-05** |
| **GO Process** | **GO:0034641** | **Cellular nitrogen compound metabolic process** | **38** | **1.69E-05** |
| GO Process | GO:0044237 | Cellular metabolic process | 57 | 1.69E-05 |
| **GO Process** | **GO:0090304** | **Nucleic acid metabolic process** | **30** | **1.69E-05** |
| **GO Process** | **GO:1901360** | **Organic cyclic compound metabolic process** | **34** | **3.80E-05** |
| GO Process | GO:0008152 | Metabolic process | 62 | 0.00021 |
| GO Process | GO:0044238 | Primary metabolic process | 56 | 0.00083 |
| GO Process | GO:0071704 | Organic substance metabolic process | 57 | 0.0022 |
| GO Process | GO:0043170 | Macromolecule metabolic process | 47 | 0.0023 |
| GO Process | GO:0009987 | Cellular process | 81 | 0.0029 |
| GO Process | GO:0006807 | Nitrogen compound metabolic process | 51 | 0.003 |
| GO Process | GO:0006259 | DNA metabolic process | 12 | 0.0056 |
| GO Process | GO:0006974 | Cellular response to DNA damage stimulus | 10 | 0.0091 |
| GO Process | GO:0010467 | Gene expression | 23 | 0.0091 |

| | | | | |
|---|---|---|---|---|
| GO Process | GO:0044260 | Cellular macromolecule metabolic process | 26 | 0.0091 |
| GO Process | GO:0006281 | DNA repair | 9 | 0.0107 |
| GO Process | GO:0016070 | RNA metabolic process | 19 | 0.0138 |
| GO Process | GO:0033554 | Cellular response to stress | 14 | 0.0424 |
| GO Component | GO:0005622 | Intracellular anatomical structure | 84 | 3.16E-07 |
| GO Component | GO:0043229 | Intracellular organelle | 73 | 6.60E-06 |
| GO Component | GO:0043227 | Membrane-bounded organelle | 68 | 1.40E-05 |
| GO Component | GO:0043231 | Intracellular membrane-bounded organelle | 66 | 1.81E-05 |
| GO Component | GO:0032991 | Protein-containing complex | 46 | 4.25E-05 |
| GO Component | GO:0005737 | Cytoplasm | 54 | 0.0187 |
| GO Component | GO:0110165 | Cellular anatomical entity | 90 | 0.0187 |
| Monarch | FBcv:0002004 | Increased mortality | 73 | 2.97E-06 |
| Monarch | FBcv:0002019 | Increased mortality during development | 69 | 5.68E-06 |
| Monarch | FBcv:0000351 | Lethal | 57 | 0.00012 |
| Monarch | FBcv:0000354 | Visible | 54 | 0.00012 |
| Monarch | FBcv:0002015 | Partially lethal | 39 | 0.0071 |
| Monarch | FBcv:0000435 | Abnormal neuroanatomy | 28 | 0.0072 |
| Monarch | FBcv:0002010 | Majority die during P-stage | 18 | 0.0072 |
| Monarch | FBcv:0002005 | Lethal - all die during P-stage | 17 | 0.0073 |
| Monarch | FBcv:0002000 | Lethal - all die before end of P-stage | 39 | 0.0143 |

| | | | | |
|---|---|---|---|---|
| Monarch | FBcv:0002020 | Some die during P-stage | 37 | 0.0143 |
| Monarch | FBcv:0002007 | Lethal - all die during larval stage | 11 | 0.0197 |
| Monarch | FBcv:0000352 | Partially lethal - majority die | 28 | 0.0441 |
| Monarch | FBcv:0002039 | Some die during pupal stage | 30 | 0.0441 |
| COMPARTMENTS | GOCC:0032991 | Protein-containing complex | 48 | 0.00011 |
| COMPARTMENTS | GOCC:0005622 | Intracellular | 62 | 0.004 |

**Table S11:** GO enrichment amount genes that moved between autosomes in Hippoboscidae.

| #category | term ID | term description | observed gene count | false discovery rate |
|---|---|---|---|---|
| GO Process | GO:0009950 | Dorsal/ventral axis specification | 6 | 0.0157 |
| UniProt Keywords | KW-0995 | Kinetochore | 3 | 0.016 |
| UniProt Keywords | KW-0720 | Serine protease | 6 | 0.0217 |
| UniProt Keywords | KW-0137 | Centromere | 3 | 0.022 |
| UniProt Keywords | KW-0645 | Protease | 8 | 0.0396 |

**Table S12:** GO enrichment amount genes that moved between autosomes in Drosophilidae.

| #category | term ID | term description | observed gene count | false discovery rate |
|---|---|---|---|---|
| GO Component | GO:0032991 | Protein-containing complex | 57 | 0.00062 |
| GO Component | GO:0043226 | Organelle | 90 | 0.00067 |
| GO Component | GO:0005622 | Intracellular anatomical structure | 98 | 0.0027 |
| GO Component | GO:0022627 | Cytosolic small ribosomal subunit | 6 | 0.0029 |
| GO Component | GO:0043229 | Intracellular organelle | 86 | 0.0029 |
| GO Component | GO:0110165 | Cellular anatomical entity | 123 | 0.0029 |
| GO Component | GO:0015935 | Small ribosomal subunit | 7 | 0.0036 |
| GO Component | GO:0043227 | Membrane-bounded organelle | 78 | 0.012 |
| GO Component | GO:0022626 | Cytosolic ribosome | 7 | 0.0161 |
| GO Component | GO:0044391 | Ribosomal subunit | 9 | 0.0172 |
| GO Component | GO:0140513 | Nuclear protein-containing complex | 22 | 0.0205 |
| Reactome | DME-72695 | Formation of the ternary complex, and subsequently, the 43S complex | 7 | 0.0023 |
| Reactome | DME-72649 | Translation initiation complex formation | 7 | 0.0036 |
| Reactome | DME-72702 | Ribosomal scanning and start codon recognition | 7 | 0.0036 |
| Reactome | DME-8953854 | Metabolism of RNA | 17 | 0.0042 |
| Reactome | DME-72689 | Formation of a pool of free 40S subunits | 7 | 0.0166 |

| | | | | |
|---|---|---|---|---|
| Reactome | DME-156827 | L13a-mediated translational silencing of Ceruloplasmin expression | 7 | 0.0333 |
| Reactome | DME-72706 | GTP hydrolysis and joining of the 60S ribosomal subunit | 6 | 0.041 |
| Reactome | DME-1799339 | SRP-dependent cotranslational protein targeting to membrane | 6 | 0.0411 |
| Reactome | DME-975956 | Nonsense Mediated Decay (NMD) independent of the Exon Junction Complex (EJC) | 6 | 0.0425 |
| Monarch | FBcv:0000443 | Minute | 6 | 0.00068 |
| COMPARTMENTS | GOCC:0015935 | Small ribosomal subunit | 7 | 0.0109 |
| COMPARTMENTS | GOCC:0022627 | Cytosolic small ribosomal subunit | 6 | 0.0109 |
| COMPARTMENTS | GOCC:0032991 | Protein-containing complex | 54 | 0.0109 |
